# Longitudinal multi-omics reveals pathogenic *TSC2* variants disrupt developmental trajectories of human cortical organoids derived from Tuberous Sclerosis Complex

**DOI:** 10.1101/2024.10.07.617121

**Authors:** Weibo Niu, Shaojun Yu, Xiangru Li, Zhen Wang, Rui Chen, Christina Michalski, Arman Jahangiri, Youssef Zohdy, Joshua J Chern, Ted J Whitworth, Jianjun Wang, Jie Xu, Ying Zhou, Zhaohui Qin, Bingshan Li, Michael J Gambello, Junmin Peng, Zhexing Wen

## Abstract

Tuberous Sclerosis Complex (TSC), an autosomal dominant condition, is caused by heterozygous mutations in either the *TSC1* or *TSC2* genes, manifesting in systemic growth of benign tumors. In addition to brain lesions, neurologic sequelae represent the greatest morbidity in TSC patients. Investigations utilizing *TSC1/2*-knockout animal or human stem cell models suggest that TSC deficiency-causing hyper-activation of mTOR signaling might precipitate anomalous neurodevelopmental processes. However, how the pathogenic variants of *TSC1/2* genes affect the longitudinal trajectory of human brain development remains largely unexplored. Here, we employed 3-dimensional cortical organoids derived from induced pluripotent stem cells (iPSCs) from TSC patients harboring *TSC2* variants, alongside organoids from age- and sex-matched healthy individuals as controls. Through comprehensively longitudinal molecular and cellular analyses of TSC organoids, we found that *TSC2* pathogenic variants dysregulate neurogenesis, synaptogenesis, and gliogenesis, particularly for reactive astrogliosis. The altered developmental trajectory of TSC organoids significantly resembles the molecular signatures of neuropsychiatric disorders, including autism spectrum disorders, epilepsy, and intellectual disability. Intriguingly, single cell transcriptomic analyses on TSC organoids revealed that *TSC2* pathogenic variants disrupt the neuron/reactive astrocyte crosstalk within the NLGN-NRXN signaling network. Furthermore, cellular and electrophysiological assessments of TSC cortical organoids, along with proteomic analyses of synaptosomes, demonstrated that the *TSC2* variants precipitate perturbations in synaptic transmission, neuronal network activity, mitochondrial translational integrity, and neurofilament formation. Notably, similar perturbations were observed in surgically resected cortical specimens from TSC patients. Collectively, our study illustrates that disease-associated *TSC2* variants disrupt the neurodevelopmental trajectories through perturbations of gene regulatory networks during early cortical development, leading to mitochondrial dysfunction, aberrant neurofilament formation, impaired synaptic formation and neuronal network activity.

## INTRODUCTION

Tuberous Sclerosis Complex (TSC) is an autosomal dominant disorder caused by heterozygous pathogenic variants in the *TSC1* or *TSC2* genes. It affects multiple organ systems including the brain, skin, kidneys, and lungs, leading to a diverse range of clinical manifestations ^1–3^. In addition to the brain lesions such as cortical tubers, subependymal nodules, and subependymal giant cell astrocytoma, TSC causes significant neurologic morbidity and mortality typically due to epilepsy, autism spectrum disorders (ASD), intellectual disability, and TSC-associated neuropsychiatric disorders ^1,4–6^. However, even patients without tubers can exhibit significant developmental deficits ^7,8^, including intellectual disability and autism, suggesting that neurological manifestations in TSC might stem from broader neurodevelopmental disruptions that extend beyond the visible lesions typically associated with the disorder and these symptoms may arise from mechanisms independent of cortical tubers.

While neurological manifestations of TSC typically emerge in late adolescence to early adulthood ^9^, it is posited that the etiology is rooted in detrimental neuropathological events occurring during the early stages of brain development ^10^. Despite significant advancements in elucidating the signaling pathways underlying TSC in animal models of *TSC1/TSC2* deficiency, the impact of disease-associated *TSC1* and *TSC2* variants on human brain development remains elusive. This gap in knowledge largely stems from the challenges in acquiring live, developing human cortical tissues for study. Human induced pluripotent stem cells (iPSCs), which capture identical risk alleles as the donor individual and have the capacity to differentiate into a myriad of human cell type, offer an unprecedented opportunity to recapitulate both normal and pathologic human development. Recent studies utilizing human iPSC models, including two-dimensional cultures of neural progenitor cells (NPCs), neurons, or astrocytes, as well as three-dimensional brain organoids, have identified certain developmental impairments linked to TSC deficiency ^2,11–18^. For instance, Eichmüller et al. identified caudal late interneuron progenitor (CLIP) cells as key contributors to cortical malformations and brain tumors in TSC ^15^. However, these studies have predominantly concentrated on singular developmental stages or exclusively on atypical cell proliferation. A more thorough assessment of the effects of pathogenic *TSC1* and *TSC2* variants on the molecular and cellular progressions of human brain development is imperative. Such research is crucial for uncovering the underlying mechanisms of neuropsychiatric symptoms in TSC and for the identification of novel therapeutic targets.

In this study, we developed human cortical organoids from patient-derived iPSCs carrying *TSC2* pathogenic variants and age- and sex-matched healthy controls, as well as three isogenic lines by introducing patient-specific *TSC2* variants into a wildtype control iPSC line. Longitudinal transcriptomic analysis of these cortical organoids, conducted at four distinct developmental stages, revealed disrupted developmental trajectories in TSC, encompassing critical processes such as neurogenesis, synaptogenesis, and gliogenesis. The divergent developmental trajectory observed in TSC organoids notably mirrors the molecular signatures characteristic of various neuropsychiatric disorders, including ASDs, epilepsy, and intellectual disability. Utilizing single-cell level analysis of cell-cell communication, we found that pathogenic *TSC2* variants disrupt intercellular interactions, specifically targeting neuron/reactive astrocyte communication within the NLGN-NRXN signaling network. Moreover, comprehensive cellular and electrophysiological evaluations of TSC cortical organoids, combined with in-depth proteomic analyses of synaptosomes, revealed that pathogenic *TSC2* variants induce disturbances in mitochondrial translational integrity, neurofilament formation, synaptic transmission, and the overall activity of neuronal networks. Notably, increased neurofilament formation, compromised mitochondrial integrity, and enhanced synapse formation were observed in cortical specimens surgically excised from TSC patients, underscoring their direct association with pathogenic *TSC2* variants. In summary, our study demonstrates that *TSC2* pathogenic variants disrupt the neurodevelopmental trajectories through perturbations of gene regulatory networks during early cortical development, leading to mitochondrial dysfunction, aberrant neurofilament formation, impaired synaptic formation, and compromised neuronal network activity.

## RESULTS

### *TSC2* pathogenic variants accelerate the development of cortical organoids

To explore the impact of disease-linked *TSC2* variants on cortical development, we established iPSCs from four TSC patients harboring *TSC2* heterozygous variants (TSC001: *TSC2* c.G4502A, p.R1501H; TSC007: *TSC2* c.116_118del, p.39_40del; TSC009: *TSC2* c.C3643T, p.R1215X; and TSC010: *TSC2* c.G4514C, p.R1505P; Supplementary Table 1), as well as four sex- and age-matched healthy controls (C1-2, 11C1, 426, and PGP1; Supplementary Table 1). Additionally, we generated one heterozygous isogenic line (named as TSC010Het) and two homozygous isogenic lines (named as TSC009HO and TSC010HO) by introducing the variants of TSC009 (*TSC2* c.C3643T, p.R1215X) and TSC010 (*TSC2* c.G4514C, p.R1505P), respectively, into a wildtype control iPSC line (PGP1) using CRISPR/Cas9 gene editing technology. All the pathogenic variants in iPSCs were validated by Sanger sequencing (Extended Data Fig. 1).

To assess the effects of pathogenic *TSC2* variants on the molecular trajectories of cortical development, we generated three-dimensional cortical organoids from these iPSC lines using established protocol ^19^ and conducted longitudinal transcriptomic analyses on both TSC and control cortical organoids across four distinct stages of development (day 35, 56, 98, and 126) (Fig. 1a). Comparison of our organoid transcriptomic data with that of human dorsolateral prefrontal cortex samples across six life stages, from fetal development to aged tissue ^20^, revealed the highest correlation with fetal brain tissues (Fig. 1b, Extended Data Fig. 2a). Interestingly, TSC organoids exhibited a developmental shift, aligning more closely with fetal brain tissues (Fig. 1b, Extended Data Fig. 2a). Further mapping our organoid transcriptomic data to the BrainSpan dataset from human dorsolateral prefrontal cortex (https://www.brainspan.org/), a benchmark in vivo reference for brain development, also confirmed that both control and TSC organoids closely resemble the molecular signatures of the human fetal cortex (Extended Data Fig. 2b-c). Consistently, TSC organoids exhibited a developmental shift from resembling first trimester fetal brain tissues to those of the second trimester (Fig. 1b, Extended Data Fig. 2b-c), indicating an accelerated development. Furthermore, dorsal forebrain-specific marker genes, including *PAX6*, *SATB2*, *BCL11B*, and *NEUROD1*, are highly expressed in both control and TSC organoids (Extended Data Fig. 2d), confirming their identity of dorsal forebrain. While the *TSC2* variants did not impact the overall size of cortical organoids (Extended Data Fig. 2e-f) or the proportion of SOX2^+^ neural progenitor cells (NPCs) among all DAPI^+^ cells in general (Extended Data Fig. 2h), we observed a significant increase in Ki67^+^ proliferating cells among all SOX2^+^ NPCs (Extended Data Fig. 2g, i) in TSC organoids, suggesting enhanced neurogenesis due to *TSC2* variants.

**Figure 1.**
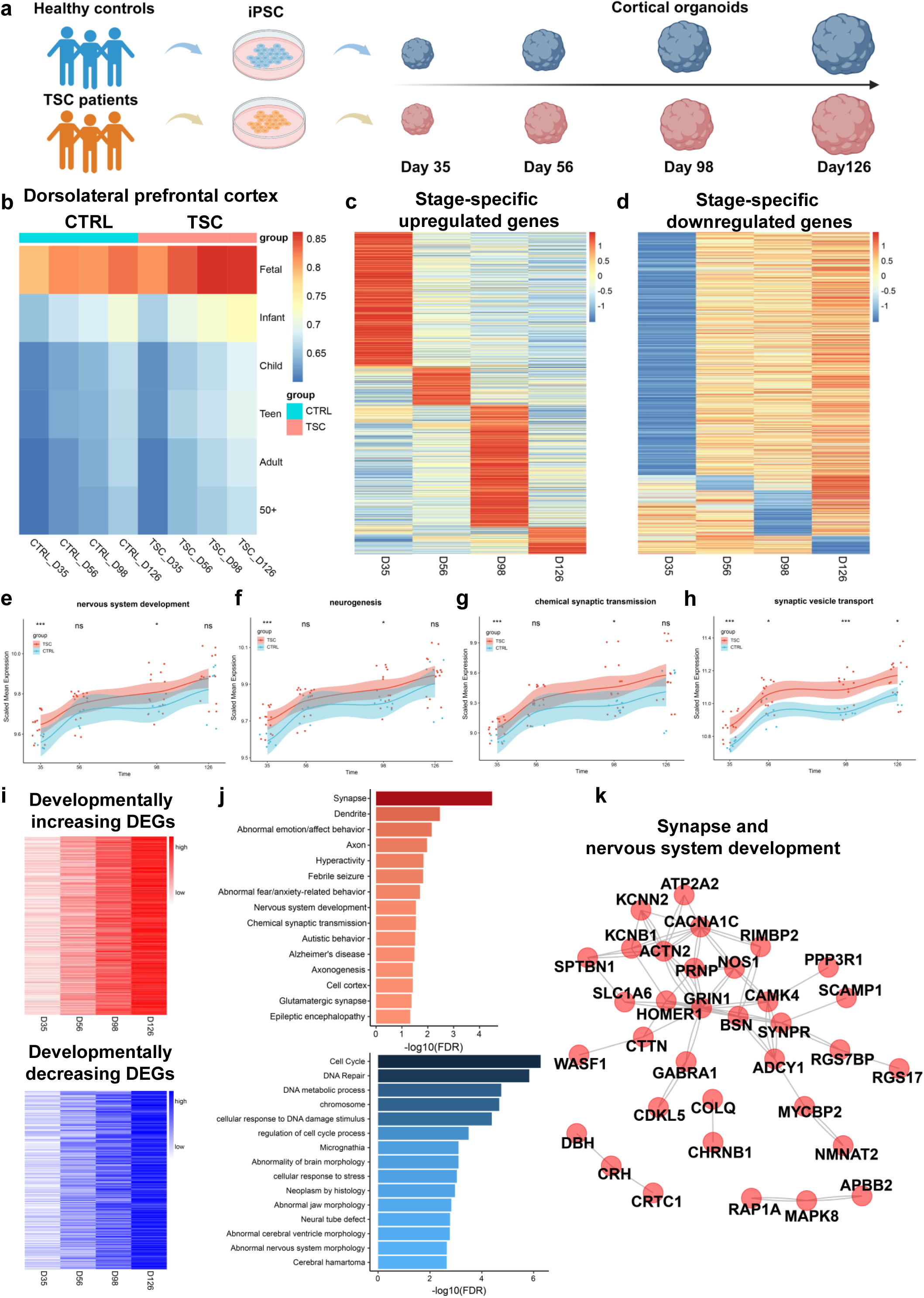
*TSC2* pathogenic variants accelerate the development of cortical organoids. **(a)** Schematic of the collection time points of cortical organoids derived from healthy controls and TSC patients for bulk RNA sequencing: day 35 (D35), day 56 (D56), day 98 (D98), and day 126 (D126). **(b)** TSC organoids exhibit accelerated development. Heatmap shows Pearson’s correlation analysis of transcriptomic datasets from day 35, 56, 98, and 126 organoids and published datasets from the human dorsolateral prefrontal cortex (DLPFC) from the BrainSpan dataset. Colors indicate the averages for biological replicates. **(c)** Heatmap of development-stage-specific upregulated differentially expressed genes (DEGs). Colors indicate normalized relative genes expression levels. **(d)** Heatmap of development-stage-specific downregulated DEGs, where the color scale represents normalized relative gene expression levels (adjusted *P*-value < 0.05, absolute log2 fold-change > 1.5). **(e-h)** Normalized mean expression of neuronal and synaptic module genes involved in nervous system development (e), neurogenesis (f), chemical synaptic transmission (g), and synaptic vesicle transport (h) in the control and TSC organoids. X-axis represents differentiation day, and Y-axis represents normalized mean expression. The red and blue area surrounding the trajectory line signifies the 95% confidence interval. \**P* < 0.05, \*\*\**P* < 0.001, one-tailed t-test. **(i)** Dynamics of DEGs during TSC organoids development. Shown are developmentally increasing DEGs (top) and developmentally decreasing DEGs (bottom) in TSC organoids during development. Colors indicate relative genes expression levels. **(j)** Pathway analyses of developmentally increasing DEGs and developmentally decreasing DEGs in TSC organoids (FDR < 0.05). **(k)** Gene interaction analysis of the synapse and nervous system development pathways reveals a highly interactive functional network of DEGs in TSC organoids. DEGs, differentially expression genes.

To further examine the intricate dynamics of gene expression throughout the development of TSC organoids, we identified differentially expressed genes (DEGs) at each stage (Fig. 1c-d; Supplementary Table 2-5), including 456, 199, 396, and 69 upregulated DEGs, as well as 887, 109, 156, and 53 downregulated DEGs at day 35, 56, 98, and 126, respectively (Extended Data Fig. 3a, Extended Data Fig. 4a, Extended Data Fig. 5a, and Extended Data Fig. 6a). The heatmaps of DEGs confirmed consistent expression patterns across each line of both control and TSC organoids (Extended Data Fig. 3b, Extended Data Fig. 4b, Extended Data Fig. 5b, and Extended Data Fig. 6b). Gene ontology (GO) analyses of DEGs at each stage revealed that upregulated DEGs in TSC organoids are enriched in nervous system development, neurogenesis, synapse, and axonogenesis (Extended Data Fig. 3d-e; Extended Data Fig. 4c-d; Extended Data Fig. 5c-d; and Extended Data Fig. 6c-d). Gene interaction analyses showed a highly interactive functional network of DEGs associated with neurogenesis and synaptic processes (Extended Data Fig. 3f-g; Extended Data Fig. 4e-f; Extended Data Fig. 5e-f). To further delineate the trajectories of specific biological processes in cortical organoids altered by *TSC2* variants, we utilized co-expression modules ^21^ from in vivo brain development to map the expression dynamics of both TSC and control organoids. We observed that several key biological modules, including nervous system development, neurogenesis, synaptic transmission and vesicle transport, were significantly upregulated in TSC organoids during development (Fig. 1e-h). Notably, we identified a set of DEGs that either developmentally increased or decreased across developmental stages (Fig. 1i). Intriguingly, the developmentally increased genes in TSC organoids were enriched in biological pathways associated with synapse, dendrite, axon, nervous system development, and chemical synaptic transmission, while developmentally decreased genes were involved in pathways linked to cell cycle, DNA repair, and chromosome (Fig. 1j). STRING analysis revealed a highly interactive network of DEGs associated with synapse and nervous system development (Fig. 1k). Our findings thus indicate that pathogenic variants in the *TSC2* gene significantly influence the developmental trajectories of cortical organoids, leading to enhanced synaptic development and accelerated maturation of the overall nervous system.

### Divergent developmental trajectory of TSC organoids significantly resemble the molecular signatures of neuropsychiatric disorders

TSC patients experience severe neurological manifestations: 90% with epilepsy, 90% with TSC-associated neuropsychiatric disorders (TAND), 50% are diagnosed with ASD, and 50% diagnosed with intellectual disability (ID) ^1^. Indeed, we identified a number of DEGs associated with neurodevelopmental and neuropsychiatric disorders that were significantly dysregulated during the development of TSC organoids (Extended Data Fig. 7). To further examine if the specific expression patterns of TSC organoids during development are associated with risk gene trajectories of neuropsychiatric disorders, we mapped genes associated with ASDs, epilepsy, ID, and schizophrenia (SCZ) onto the gene expression data of TSC organoids (Fig. 2 and Extended Data Fig. 8) ^21^. We conducted a weighted gene co-expression network analysis (WGCNA) and analyzed genes linked to disorders based on their temporal expression patterns in both control and TSC organoids. These analyses identified distinct clusters that represent unique temporal trajectories for each disorder. Further annotation using gene ontology highlighted that these clusters correspond to specific developmental trajectories and distinct biological processes. We found that ASD risk genes clustered into five developmental trajectories (Fig. 2a), epilepsy genes formed three clusters (Fig. 2d), while both ID and SCZ genes formed four clusters each (Fig. 2g and Extended Data Fig. 8a). We next identified two trajectory patterns shared across neurodevelopmental and neuropsychiatric disorders. The first pattern, observed in various modules including ASD-ME1, ASD-ME2, epilepsy-ME2, ID-ME2, and SCZ-ME3, displayed an increasing trend during differentiation (Fig. 2b, e, h and Extended Data Fig. 8b). These clusters revealed notable differences in the expression of risk gene subsets between control and TSC organoids and were particularly enriched in biological processes such as neurogenesis, synapse, seizure, neurodevelopmental abnormality, as well as glial cells (Fig. 2c, f, i and Extended Data Fig. 8c). Moreover, the majority of epilepsy-related genes were found clustered into epilepsy-ME2, which is associated with synaptic signaling, ion channel activity, and includes a range of excitatory and inhibitory neuronal genes such as *SYN1*, *MEF2C*, *RELN*, *SLC1A2*, *SCN9A*, *KCNQ2*, *KCNQ3*, *GRIN2B*, and *GABBR2*. Intriguingly, many DEGs in TSC organoids are ASD susceptibility genes from the SFARI Gene database (https://gene.sfari.org/) (Extended Data Fig. 3c), highlighting shared molecular pathways that may contribute to the co-occurrence of TSC and ASD. Furthermore, DisGeNET analysis ^22^ found that upregulated DEGs in TSC organoids are predominantly enriched in disease pathways associated with various types of seizures (Extended Data Fig. 9), including epileptic seizures, which aligns with our pathway analyses of disease risk gene trajectories, consistently pointing to a significant relationship between the molecular alterations in TSC and the prevalence of seizure disorders. On the other hand, a second pattern was identified in the modules epilepsy-ME1, ID-ME1, and SCZ-ME4 (Fig. 2b, e, h and Extended Data Fig. 8b), which exhibited a decreasing trend during differentiation. These clusters demonstrated significant expression pattern differences between control and TSC organoids and were enriched in processes related to seizure activity and abnormal nervous system physiology. Overall, these findings suggest significant divergences in the expression trajectories of genes associated with neurodevelopmental and neuropsychiatric disorders between control and TSC organoids.

**Figure 2.**
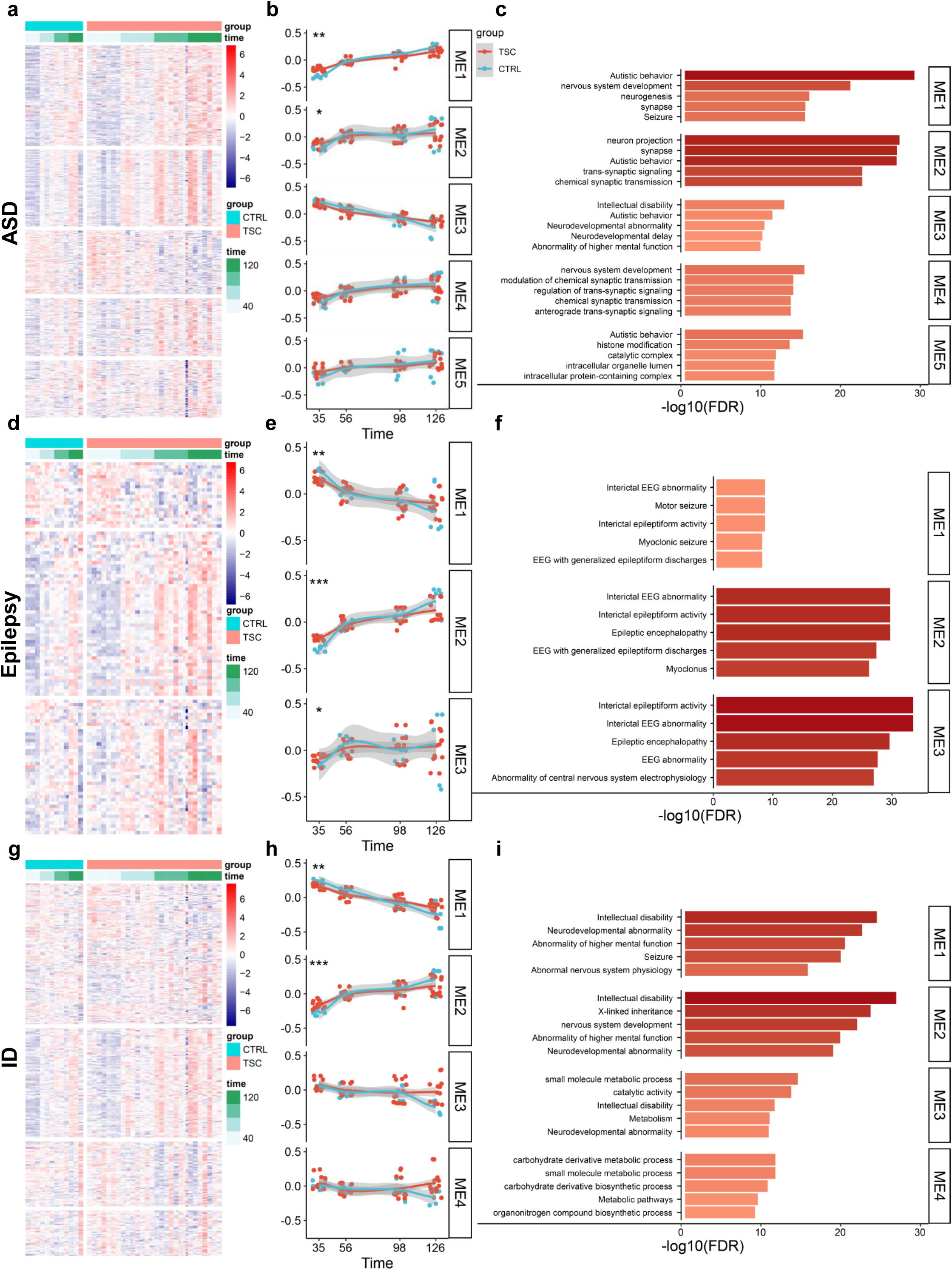
Divergent developmental trajectory of TSC organoids significantly resemble the molecular signatures of neuropsychiatric disorders. Mapping of genes associated with ASD **(a-c)**, epilepsy **(d-f)**, and ID **(g-i)** onto module trajectories shows significant differences between TSC and control organoids. The initial column displays the grouping of standardized normalized expressions of genes linked to a disorder **(a, d, g)**. The genes, arranged in rows, are grouped through hierarchical clustering based on the Euclidean distance between them. The samples, organized in columns, are sorted by the day of differentiation (indicated by green bars), starting with the earliest 35 days on the left and progressing to the latest timepoints 126 days on the right. The middle column **(b, e, h)** displays the first principal component of the eigengenes for the identified gene clusters. The grey area surrounding the trajectory line signifies the 95% confidence interval. \**FDR* < 0.05, \*\**FDR* < 0.01, \*\*\**FDR* < 0.001, two-sided Mann-Whitney test. The last column **(c, f, i)** presents the top GO terms that are enriched in the identified clusters.

### *TSC2* variants disturb the developmental trajectories of specific cell types

The gliogenic switch occurs around day 90 in cortical organoids, marking the onset of astrogenesis^23^. To further elucidate the impact of pathogenic *TSC2* variants on specific cell types at the single cell resolution, we performed single-cell RNA sequencing (scRNA-seq) on TSC and control cortical organoids at day 98. Voxhunt analysis ^24^ confirmed that our cortical organoids exhibited a high representation of the dorsal pallium (Extended Data Fig. 10a), indicating that the organoids closely mimic this region of the human cortex. Moreover, the unsupervised clustering in Uniform Manifold Approximation and Projection (UMAP) plots demonstrated the reproducibility across different cell lines and experimental batches (Extended Data Fig. 10b). We identified a total of 13 clusters: immature astrocytes (cluster 0), glial progenitor cells (cluster 1), deep layer excitatory neurons (cluster 2), astrocytes (cluster 3), upper layer excitatory neurons (cluster 4), mature interneurons (cluster 5), intermediate progenitor cells (cluster 6), immature preoptic area (pOA) interneurons (cluster 7), reactive astrocytes (cluster 8), immature medial ganglionic eminence (MGE) interneurons (cluster 9), proliferating neural progenitor cells (cluster 10), radial glial cells (cluster 11), and outer radial glia cells (cluster 12), which were systematically classified as representing neural lineages based on the cell type specific molecular signatures in combination with gene ontology (GO) analysis ^25,26^ (Fig. 3a and Extended Data Fig. 10c). To ascertain whether *TSC2* variants influence the cellular composition, we quantified the proportion of each cell cluster present within the organoids (Fig. 3b and Extended Data Fig. 10d). We found that the numbers of immature astrocytes, deep layer excitatory neurons, upper layer excitatory neurons, mature interneurons, reactive astrocytes, and proliferating neural progenitor cells (clusters 0, 2, 4, 5, 8, and 10, respectively) were increased in TSC organoids (Fig. 3b and Extended Data Fig. 10d). Indeed, we observed increased expression of *BCL11B* (CTIP2) in the deep layer excitatory neurons (Extended Data Fig. 12a). Immunostaining confirmed that the percentage of CTIP2^+^ deep layer excitatory neurons was significantly increased in TSC organoids (Fig. 3c-d). On the other hand, we observed a downregulation of glial progenitor cells and immature pOA interneurons (clusters 1 and 7) in TSC organoids (Fig. 3b and Extended Data Fig. 10d). To further explore the developmental trajectories of these cell populations, we conducted RNA velocity analyses on both control and TSC cortical organoids (Fig. 3e). Starting from the radial glia cell population (cluster 11), we identified two distinct trajectories: one leading towards an increase in immature astrocytes (cluster 0) and particularly reactive astrocytes (cluster 8) in TSC organoids; the other towards an increase in deep layer excitatory neurons (cluster 2) and upper layer excitatory neurons (cluster 4) in TSC organoids. Our data thus further delineated the altered developmental trajectories of TSC organoids at the single-cell resolution.

**Figure 3.**
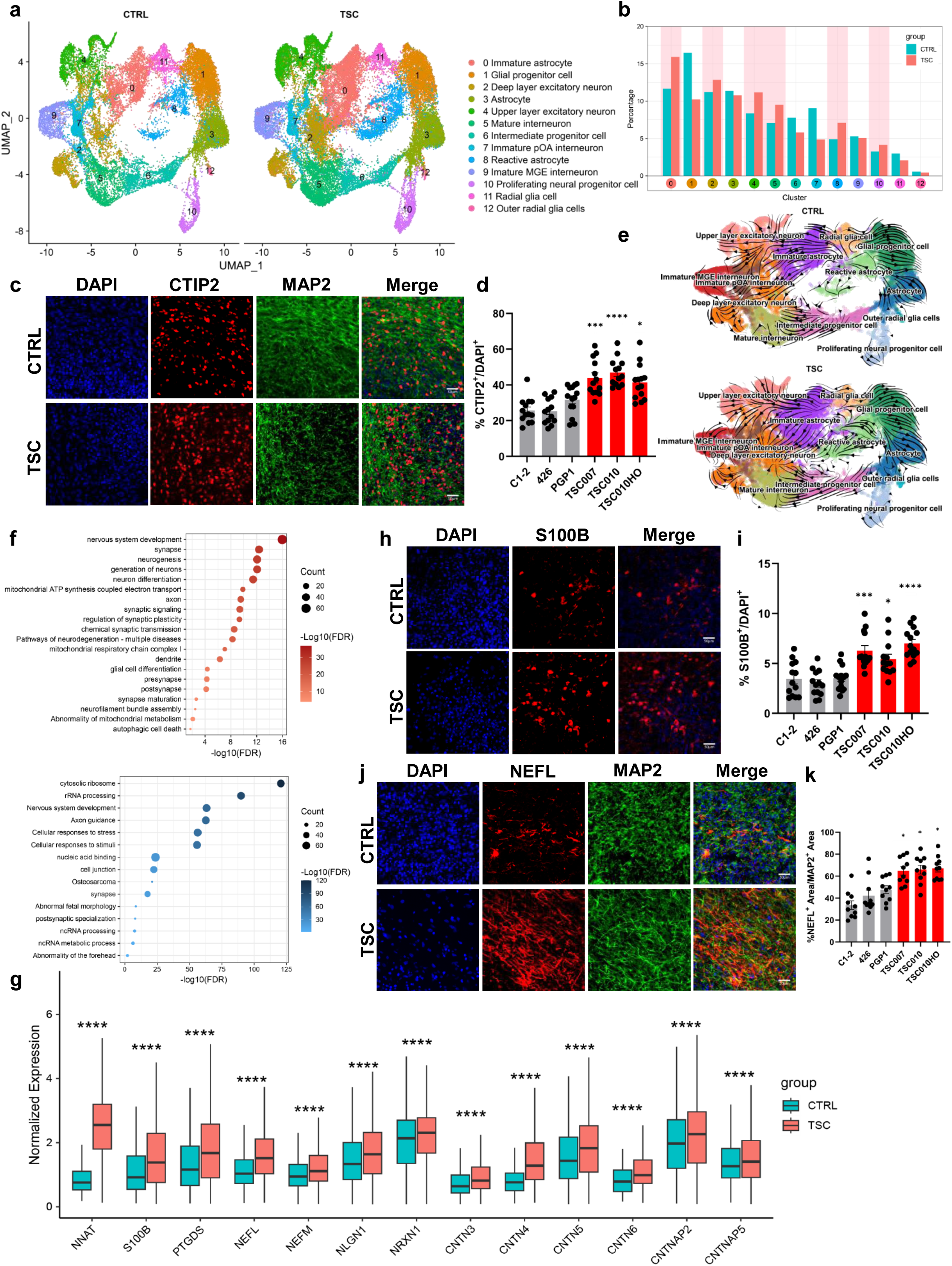
*TSC2* variants disturb the developmental trajectories of specific cell types. **(a)** UMAP projection of 98-day-old control (left) and TSC (right) cortical organoids reveals differential changes in the populations of specific cell types in TSC compared to the control. Within individual clusters, distinct neural cell types were identified and labeled. **(b)** Quantification of cell type composition of 13 clusters illustrated in (a) in control and TSC organoids. **(c-d)** Increased differentiation of cortical neurons in TSC organoids. Shown are representative images (c) and stereological quantification (d) of the proportion of CTIP2^+^ cortical neurons within DAPI^+^ cells in both control and TSC organoids at Day 98. Scale bars, 50 μm. Data are presented as mean ± s.e.m. (n = 13-14 for each line from three independent experiments. \**P* < 0.05, \*\*\**P* < 0.001, \*\*\*\**P* < 0.0001, one-way ANOVA). **(e)** RNA velocity of 98-day-old control (top) and TSC (bottom) cortical organoids. The annotation of the cluster corresponds to (a). From the radial glia cell population (cluster 11), two trajectories emerge: one trajectory leads toward immature astrocytes (cluster 0) and reactive astrocytes (cluster 8), while another trajectory leads toward deep layer excitatory neurons (cluster 2) and upper layer excitatory neurons (cluster 4). **(f)** Pathway analyses of overall upregulated and downregulated DEGs in TSC organoids single-cell transcriptomic analysis. Red bubbles represent GO terms associated with upregulated genes (adjusted *P*-value < 0.05, average log2 fold-change > 0.25), while blue bubbles represent GO terms associated with downregulated genes (adjusted *P*-value < 0.05, average log2 fold-change < -0.25). **(g)** Normalized expressions of DEGs in TSC organoids, including early-stage astrocyte marker, NNAT; mature astrocyte marker, S100B; caudal late interneuron progenitor (CLIP) cell marker, PTGDS; neurofilament light chain, NEFL; neurofilament medium chain, NEFM; adhesion molecules and autism-risk genes, including *NLGN1*, *NRXN1*, *CNTN3*, *CNTN4*, *CNTN5*, *CNTN6*, *CNTNAP2*, *CNTNAP5* (\*\*\*\**P* < 0.0001, t test,). (**h-i**) Increased gliogenesis in TSC organoids. Shown are representative images (h) and stereological quantification (i) of the proportion of S100B^+^ astrocytes within DAPI^+^ cells, in both control and TSC organoids at Day 98, scale bars, 50 μm. Data are presented as mean ± s.e.m. (n = 13-14 for each line from three independent experiments, \**P* < 0.05, \*\*\**P* < 0.001, \*\*\*\**P* < 0.0001, one-way ANOVA). **(j-k)** Elevated NEFL level in TSC organoids. Shown are representative images (j) and stereological quantification (k) of the percentage of NEFL^+^ area relative to MAP2^+^ area in control and TSC organoids. Data are presented as mean ± s.e.m. (n = 10 for each line from three independent experiments, \**P* < 0.05, one-way ANOVA), scale bars, 50 μm.

Pathway analyses on the overall DEGs from single-cell transcriptomes (Supplementary Table 6) revealed that upregulated DEGs in TSC organoids are enriched in biological pathways associated with synapse, neurogenesis, neuron differentiation, axon, glial cell differentiation, abnormality of mitochondrial metabolism, and neurofilament bundle assembly (Fig. 3f). We further investigated the specific DEGs within each cell cluster to gain a more nuanced understanding of the cellular dynamics and molecular alterations occurring in distinct cell types within the organoids (Extended Data Fig. 10e; Supplementary Table 7). Among the 13 clusters, radial glial cells (cluster 11) displayed the highest number of DEGs (Extended Data Fig. 10f). Pathway analysis demonstrated that upregulated DEGs in both immature and reactive astrocytes are enriched in biological pathways associated with synapse, neurogenesis, neuron differentiation, synaptic signaling, mitochondrial ATP synthesis coupled electron transport, as well as axon development (Extended Data Fig. 11a, i). Similarly, the analysis of upregulated DEGs in deep-layer excitatory neurons showed enrichment in terms related to neuron differentiation, neurogenesis, axonal development, synaptic function, and mitochondrial localization (Extended Data Fig. 11c). In addition, we observed that TSC organoids express a high proportion of the early-stage astrocyte marker *NNAT*, as well as the mature astrocyte marker *S100B* (Fig. 3g and Extended Data Fig. 13 a-c), consistent with our bulk transcriptomics and immunocytochemistry data (Fig. 3h-i and Extended Data Fig. 12b-c). In TSC organoids, there was a marked increase in prostaglandin D2 synthase (*PTGDS*; Fig. 3g and Extended Data Fig. 12d). PTGDS, a caudal late interneuron progenitor (CLIP) cell marker, has been shown to be highly expressed in *TSC2* mutant organoids ^15^. We also identified changes in the expression of several key genes associated with neurodevelopmental disorders in TSC organoids compared to controls, including neuronal immunoglobulin cell adhesion molecules (IgCAMs) genes *CHL1*, *CNTN4*, and *CNTN6*, glutamate receptor gene *GRIN2B*, GABA receptor genes *GABBR2*, and solute carrier family gene *SLC1A3* (Extended Data Fig. 12). CHL1, CNTN6, and CNTN4 have been shown to play a key role in regulating neurogenesis and synaptic function, which may confer a greater risk of ASD development ^27^. Additionally, research indicates that neuronal IgCAMs share similar interaction networks and are involved in common signaling pathways crucial for axon guidance ^27^. Simultaneous dysregulation of several contactins and other IgCAMs in cortical tubers could intensify the observed pathological phenotypes in TSC ^28^.

Another hallmark we identified in TSC organoids is the upregulation of neurofilament genes, including *NEFL* and *NEFM* (Fig. 3g and Extended Data Fig. 12 e-f). In central nervous system (CNS), neurofilaments (NFs) are structured as heteropolymers comprising four subunits: neurofilament light chain (NEFL), neurofilament medium chain (NEFM), neurofilament heavy chain (NEFH), and either α-internexin or peripherin ^29–31^. An increase in neurofilament medium and heavy chains has been described in several neurological disorders, such as Alzheimer’s disease, multiple sclerosis, and amyotrophic lateral sclerosis ^32–34^. In TSC cortical organoids, *NEFL* and *NEFM* exhibited higher expression levels in several cell types, including upper-layer and deep layer excitatory neurons as well as interneurons (Extended Data Fig. 14a-b). To confirm the increase of neurofilaments in TSC organoids, we performed NEFL and NEFM immunostaining (Fig. 3j and Extended Data Fig. 13d). Quantitative analysis revealed that the percentage of total NEFL^+^ and NEFM^+^ area relative to the entire MAP2^+^ area in TSC organoids was significantly increased compared to controls (Fig. 3k and Extended Data Fig. 13e).

### Pathogenic *TSC2* variants disrupt cell-cell communication through alterations in the NLGN-NRXN signaling network

Through single-cell transcriptomic analyses, we also identified a set of genes encoding adhesion molecules are significantly upregulated in TSC organoids, including postsynaptic cell adhesion molecule neuroligin 1 (*NLGN1*), its presynaptic partner neurexin 1 (*NRXN1*), contactin family genes *CNTN3*, *CNTN4*, *CNTN5*, *CNTN6*, as well as contactin associated proteins *CNTNAP2* and *CNTNAP5* (Fig. 3g and Extended Data Fig. 12g-n). Notably, all these genes are recognized as autism-risk genes that contribute to the structural stability of neurons ^35^. They regulate neurite outgrowth, synapse and spine formation, with NLGN and NRXN also modulating synaptic plasticity ^35^. To examine the expression and interactions of these genes, we created heatmaps and conducted gene interactions analyses (Fig. 4a-b). Indeed, *NLGN1* showed increased expression in immature astrocytes, deep layer excitatory neurons, and reactive astrocytes, while *NRXN1* was upregulated in immature astrocytes, intermediate progenitor cells, reactive astrocytes, and radial glia cells (Fig. 4a). The ligand-receptor pair NLGN1-NRXN1 signaling network makes the most significant contribution to the NLGN-NRXN signaling network (Extended Data Fig. 15a-b). These findings suggest that *TSC2* variants may disturb cell-cell communication in TSC organoids through ligand-receptor signaling networks, including NLGN1-NRXN1, NLGN1-NRXN3, NLGN1-CNTNAP2, and NLGN1-CNTN4.

**Figure 4.**
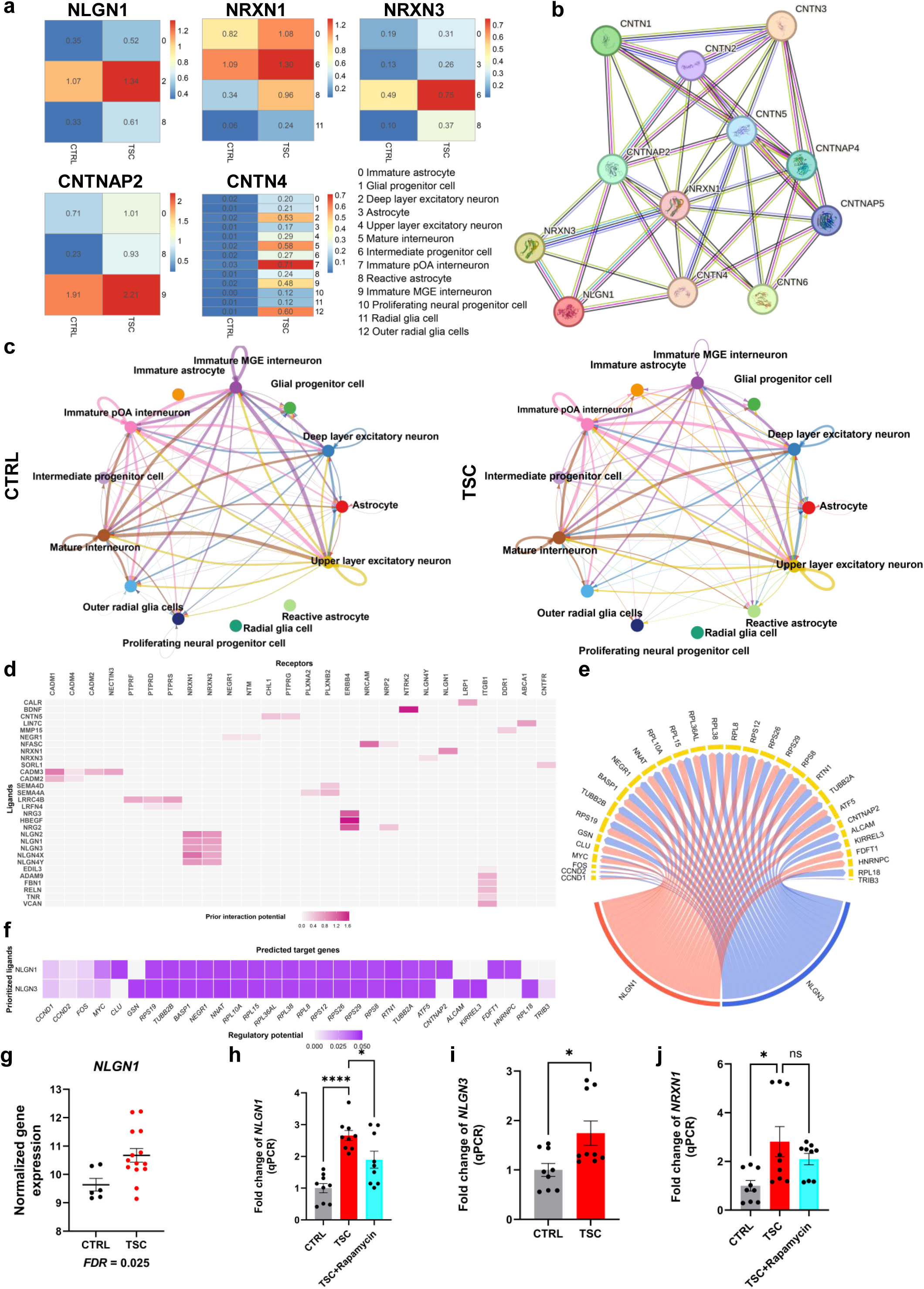
Pathogenic *TSC2* variants disrupt cell-cell communication networks. **(a)** Heatmaps show the average expressions of *NLGN1*, *NRXN1*, *NRXN3*, *CNTNAP2*, and *CNTN4* in control and TSC cortical organoids (adjusted *P*-value < 0.05, average log2 fold-change > 0). MGE, medial ganglionic eminence. pOA, preoptic area. **(b)** Gene interaction analyses of the 12 upregulated DEGs include *NLGN*, *NRXN* family genes, and *CNTN* family genes in TSC organoids (adjusted *P*-value < 0.05, average log2 fold-change > 0). **(c)** Circle plots showcase the inferred network of the NLGN-NRXN signaling pathway in control (left) and TSC (right) organoids. The thickness of all lines connecting the cell clusters indicates the strength of the interaction, as analyzed using CellChat. **(d-f)** NicheNet analysis of neuron-astrocyte interaction. Shown in (d) is the outcome of NicheNet’s ligand-receptors pairs regulating genes differentially expressed by the reactive astrocytes between TSC and controls. The top 30 prioritized ligands are expressed by deep layer excitatory neurons and upper layer excitatory neurons in TSC; shown in (e) is the NLGN1 and NLGN3 ligand-target matrix between the excitatory neuron ligands expressed in TSC, and shown in (f) are the predicted target genes expressed by the reactive astrocytes, where the matrix is colored according to the regulatory potential values. **(g)** mRNA expression level of *NLGN1* in control and TSC organoids (DESeq2, *P*-value). **(h-j)** qPCR of *NLGN1*, *NLGN3*, and *NRXN1* mRNA expression in control and TSC organoids. Data are presented as mean ± s.e.m. (n = 3 CTRL, 3 TSC, each cell line with triplicates; \**P* < 0.05, \*\*\*\**P* < 0.0001, unpaired t test).

Given that NLGN and NRXN families are integral synaptic cell adhesion molecules which not only bridge pre- and post-synaptic neurons at synapses but also regulate astrocyte-neuron interactions which orchestrate functional synapse assembly ^36–40^, we delved into the NLGN-NRXN signaling network in cell-cell communication of TSC organoids across all 13 cell clusters employing the CellChat package ^41^ (Fig. 4c, Extended Data Fig. 15). Our analyses revealed that reactive astrocytes in TSC organoids interact with different types of neurons, including deep layer excitatory neurons, upper layer excitatory neurons, mature interneurons, immature pOA interneurons, immature MGE interneurons, through the NLGN-NRXN signaling network, while these interactions are barely detectable in control organoids (Fig. 4c and Extended Data Fig. 15c-d). Our data suggests that *TSC2* variants may disturb the specific neuron/reactive astrocyte communication through alterations in the NLGN-NRXN signaling network.

To further verify the neuron/reactive astrocyte interaction, we employed NicheNet ^42^, an algorithm designed to deduce active ligand-target connections among interacting cells. We predicted networks of the ligand-receptor and ligand-target gene links among upper and deep layer excitatory neurons and reactive astrocytes. In order to predict which ligands from excitatory neurons most likely affected the gene expression in reactive astrocyte, for each couple of sender-receiver cells we obtained a list of the top 30 prioritized excitatory neuron ligands, including NLGN1, NLGN2, and NLGN3, their binding partners are NRXN1 and NRXN3 (Fig. 4d). Importantly, molecular ligand-target gene interactions, specifically related to NLGN1 communication, were closely linked to reactive astrocytes. These NLGN1 target genes include transcription factors (TFs) *ATF5*, *FOS*, and *MYC*; ribosomal protein gene *RPL10A*, *RPL15*, *RPL36AL*, *RPL38*, *RPL8*, *RPS12*, *RPS19*, *RPS26*, *RPS29*, and *RPS8*; cyclins *CCND1* and *CCND2*; tubulin beta *TUBB2A* and *TUBB2B*; neuronal growth regulator *NEGR1*; other genes such as *CNTNAP2*, *NNAT*, *BASP1*, *CLU*, *FDFT1*, *HNRNPC*, and *RTN1* (Fig. 4e-f). Remarkably, these NLGN1 target genes are pivotal in cortical development while all of them are cell type specific DEGs that are found in TSC organoids.

Additionally, the transcriptome or quantitative polymerase chain reaction (qPCR) analyses validated the increase in *NLGN1, NLGN3* and *NRXN1* in TSC organoids (Fig. 4g-j). Interestingly, the altered expression of *NLGN1* in TSC organoids could be rescued by treatment with rapamycin (20 nM) (Fig. 4h), suggesting that the disruptions in the NLGN-NRXN signaling network in TSC organoids are mTOR-dependent. In summary, our findings indicate that *TSC2* variants disrupt the cell-cell communication network between neurons and reactive astrocytes, potentially via NLGN1-dependent mechanisms.

### Altered synaptic machinery and mitochondrial translational integrity in TSC synapses

Neuron-astrocyte interaction and NLGN signaling are pivotal in synapse formation and function. Notably, our GO analyses also revealed that DEGs in TSC organoids are predominantly enriched in biological pathways associated with synapse (Fig. 1j), highlighting the synaptic dysfunction in TSC organoids. To further investigate the impact of pathogenic *TSC2* variants on synaptic machinery and assembly, we conducted deep proteomic profiling of synaptosomes isolated from control and TSC organoids at day 98. We identified a total of 6,994 unique proteins in TSC synaptosomes (Extended Data Fig. 16a), among which 561 were defined as differentially expressed proteins (DEPs) with cut off FDR < 0.05 and absolute Log2FC > 0.3, comprising 232 upregulated and 329 downregulated proteins (Extended Data Fig. 16b; Supplementary Table 8). The heatmap of these DEPs presented consistent and reproducible patterns across all controls and TSC synaptosomes (Extended Data Fig. 16c). To identify the enrichments of DEPs, we first annotated our datasets to synaptic cellular components and biological processes using the SYNGO knowledgebase ^43^ (Fig. 5a-d). The 561 DEPs were significantly enriched in GO cellular components terms presynaptic and postsynaptic annotated proteins (presynapse, GO:0098793, FDR = 2.30E-05; postsynapse, GO:0098794, FDR = 5.55E-9) (Fig. 5a-b). Among the GO biological processes terms, DEPs were significantly enriched in annotated proteins of synaptic signaling (GO:0099537, FDR = 7.95E-4), process in the presynapse (GO:0099504, FDR = 0.0221), and synapse organization (GO:0050808, FDR = 2.87E-11) (Fig. 5c-d). Subsequent pathway analyses of these DEPs by STRINGdb ^44^ (Fig. 5e-f) uncovered pathways significantly upregulated in relation to synaptic functions, including glutamatergic synapse, postsynaptic density, trans-synaptic signaling, regulation of synaptic plasticity, protein interactions at synapses, and presynaptic activities (Fig. 5e). These findings align well with our transcriptomic results. On the other hand, the downregulated DEPs, as shown by GO analyses, were predominantly enriched in mitochondrial functions including the mitochondrial ribosome, translation, and envelope (Fig. 5f). Similar results were also corroborated using the ClueGO gene ontology analysis tool ^45^. The upregulated DEPs were associated with synaptic functions (Fig. 5g), including modulation of chemical synaptic transmission, regulation of synapse organization, regulation of synapse structure or activity, and receptor localization to the synapse. Conversely, the downregulated DEPs were primarily related to mitochondrial translation (Fig. 5h). We further performed gene interaction analysis of DEPs in TSC synaptosomes and identified increased expression levels of multiple proteins on neurogenesis and trans-synaptic signaling (Fig. 5i-j), including CHL1, CNTN4, CNTNAP2, CNTNAP4, MAP2, MECP2, GRIA3, GRID1, GRM7, and SLC1A3. Decreased expression levels of multiple proteins on mitochondrion were also found (Fig. 5k), such as MRPL32, MRPL20, MRPL23, MRPS10, MRPL30, MRPL3, MRPL14, MRPL28, MRPS12, MRPS18C, MRPL24, and MRPS15. Indeed, we conducted transmission electron microscope (TEM) imaging on TSC and control organoids at day 98 and observed enlarged and autophagic mitochondria in TSC organoids (Fig. 5l). Enlarged mitochondria were significantly more prevalent in TSC organoids (Fig. 5m). Several mitochondria also possessed large granular inclusions within the matrix (Fig. 5l, middle). In addition, the proteomic analyses validated the increase in the NLGN family, including NLGN1, NLGN2, and NLGN3, in TSC synaptosomes (Fig. 5n-p), which is consistent with our transcriptome results. These findings suggest that *TSC2* variants impair the functional synapse assembly and mitochondrial integrity.

**Figure 5.**
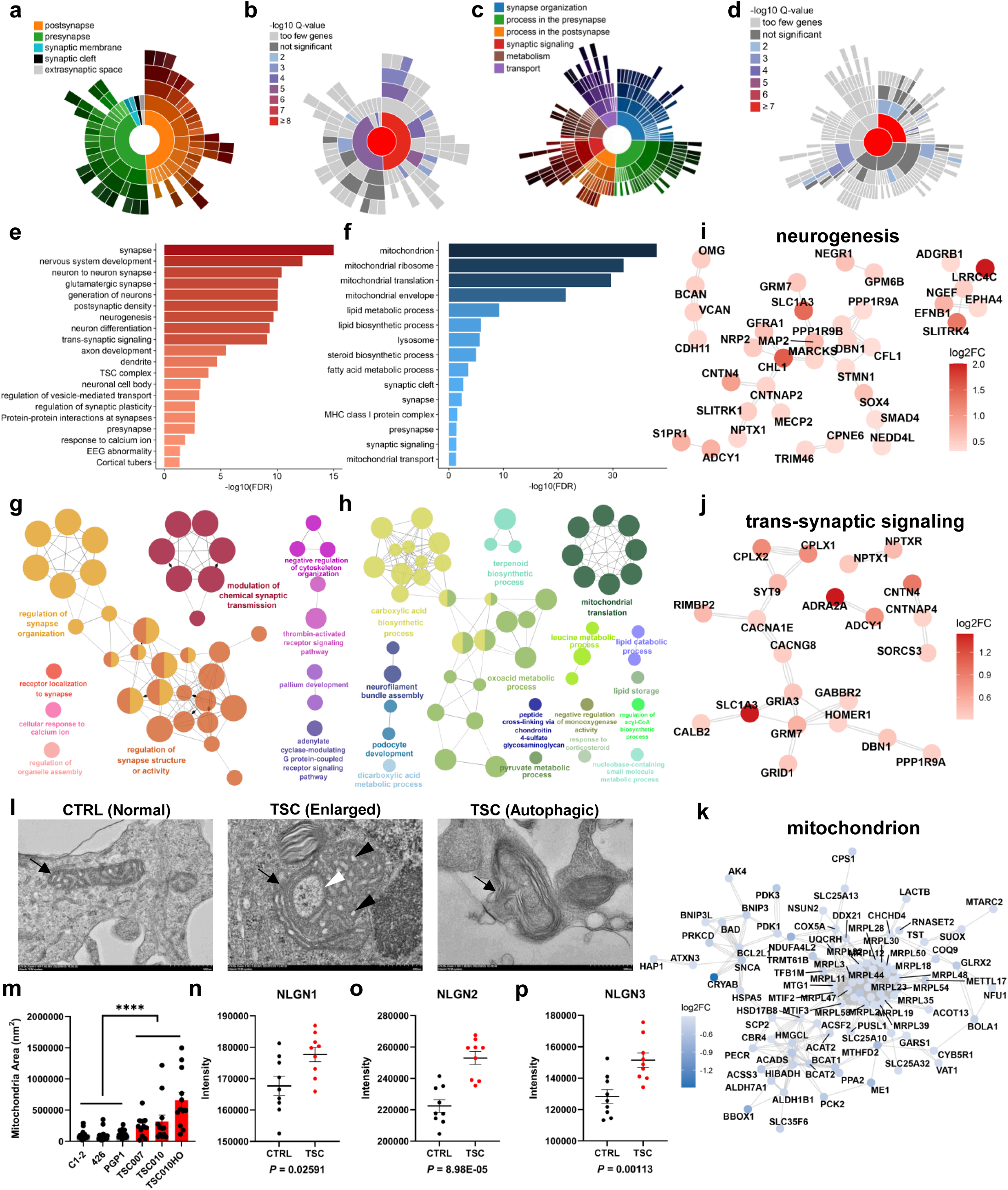
Altered synaptic machinery and mitochondrial translational integrity in TSC synapses. (a-d) Synaptic ontology analysis of 561 synaptosomes DEPs from TSC organoids compared to controls using the SYNGO tool. Shown in (a-b) are sunburst plots representing synaptic annotated GO cellular component ontologies; shown in (c-d) are sunburst plots representing synaptic annotated GO biological processes ontologies. Colors represent -log10 FDR. **(e-f)** Pathway analyses of upregulated (e) and downregulated (f) DEPs in TSC organoids (FDR < 0.05). **(g-h)** ClueGO analyses of upregulated (g) and downregulated (h) DEPs in TSC organoids (*P*-value < 0.05). Node size represents the number of mapped genes. **(i-k)** Gene interaction analyses of DEPs involved in neurogenesis pathway (i), trans-synaptic signaling pathway (j), and mitochondrion pathway (k). Circle colors indicate log2 fold-change. **(l)** Ultrastructure of mitochondria in control and TSC organoids. Mitochondrion (arrow) in the cytoplasm of a normal neuron (left); enlarged/prismatic mitochondrion (arrow) in a TSC neuron (middle) demonstrating small, membrane-bound, round to triangular electron-lucent regions in the matrix (black arrowheads), and a larger membrane-bound vesicle containing granular/fibrillar material (white arrowhead); mitochondrion (arrow) in a TSC neuron enveloped in an autophagic vacuole (right). scale bars, 500 nm. **(m)** Quantification of mitochondrial size in control and TSC organoids. Data are presented as mean ± s.e.m. (n = 19-20 mitochondria from each control cell line and n = 11-12 from each TSC cell line were analyzed, \*\*\*\**P* < 0.0001, unpaired t test). **(n-p)** The proteomic intensity of NLGN1 (n), NLGN2 (o), NLGN3 (p) in synaptosomes between control and TSC organoids. Data are presented as mean ± s.e.m. (n = 3 CTRL, 3 TSC, each cell line with triplicates; moderate t test). GO, gene ontology. DEPs, differentially expression proteins.

### *TSC2* variants increase synapse formation and enhance neuronal network activity

Considering the observed alterations in synaptic machinery and mitochondrial translation in TSC synaptosomes, we next aimed to investigate whether synapse formation is impacted by *TSC2* variants. We analyzed the density of synaptic puncta using immunocytochemistry targeting presynaptic vesicle protein Synapsin I (SYN1), postsynaptic density protein 95 (PSD95), and dendritic marker microtubule-associated protein 2 (MAP2) (Fig. 6a). The densities of SYN1^+^ and PSD95^+^ synaptic puncta were both significantly increased in TSC neurons compared to controls in day 98 cortical organoids (Fig. 6b-c), suggesting an increased formation of excitatory synapses in TSC organoids.

**Figure 6.**
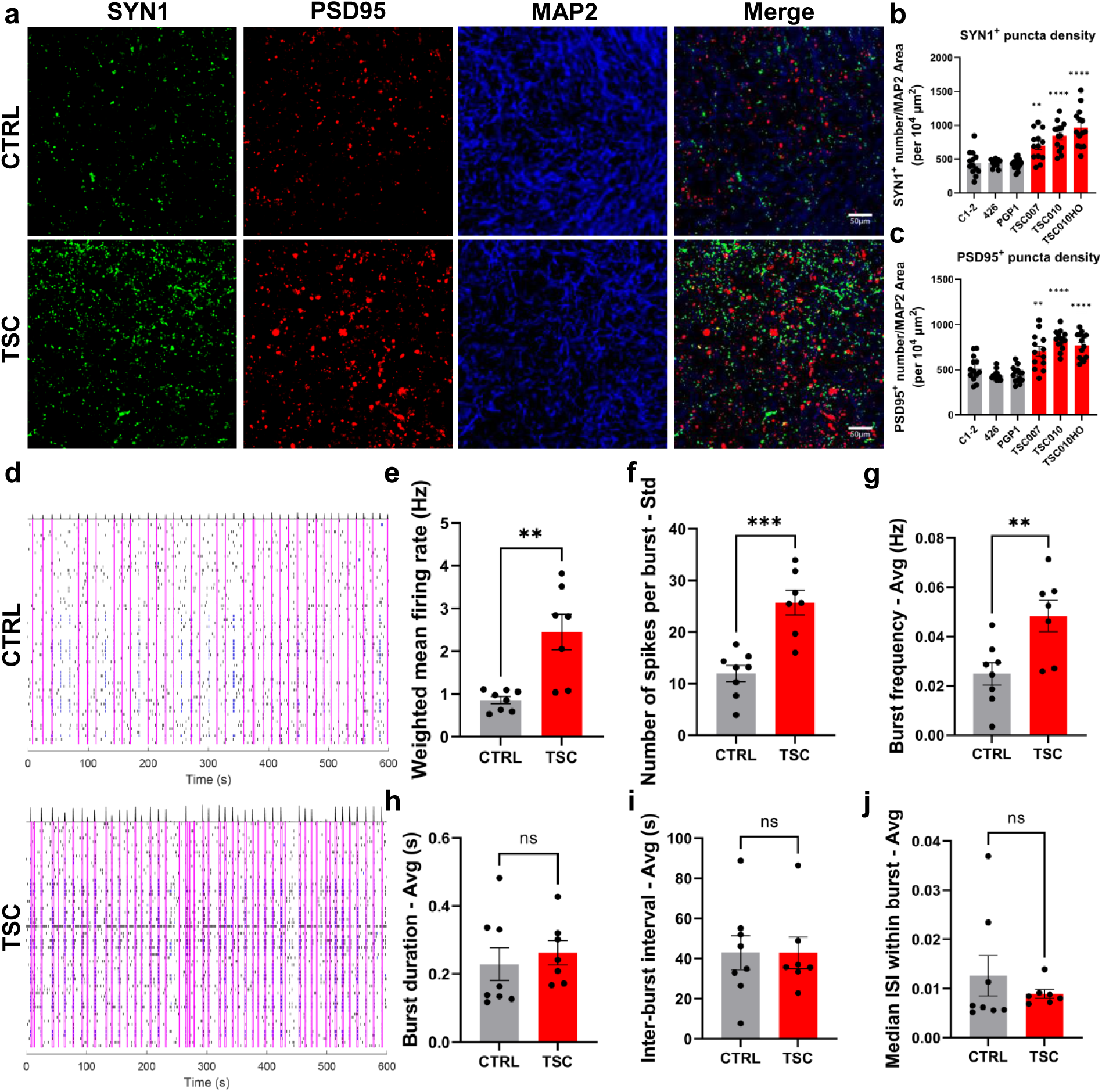
*TSC2* variants increase synapse formation and enhance neuronal network activity. (a-c) *TSC2* variants accelerate synapse formation. Shown are sample images (a) and quantification of SYN1^+^ (b) and PSD95^+^ (c) puncta density in both control and TSC organoids at Day 98, scale bars, 50 μm. Data are presented as mean ± s.e.m. (n = 13-14 for each line from three independent experiments, \*\**P* < 0.01, \*\*\*\**P* < 0.0001, one-way ANOVA). **(d-j)** TSC organoids exhibit enhanced neuronal network activity. **(d)** Raster plot of network spiking activity of control and TSC organoids, and parameters of weighted mean firing rate **(e)**, number of spikes per burst - Std **(f)**, burst frequency - Avg **(g)**, burst duration – Avg **(h)**, inter-burst interval - Avg **(i)**, and median ISI within burst – Avg **(j)**. Data are presented as mean ± s.e.m. (n = 8 for three control, n = 7 for three TSC, \*\**P* < 0.01, \*\*\**P* < 0.001, unpaired t-test). ISI, inter-spike interval.

To further examine the functional consequences of *TSC2* variants on neuronal network activity, we conducted multielectrode arrays (MEA) recordings on control and TSC cortical organoids (Fig. 6d-j). We noted a significant increase in the weighted mean firing rate (Fig. 6e), number of spikes per burst (Fig. 6f), and average burst frequency (Fig. 6g) in TSC organoids compared to the controls. However, there were no significant differences in the average burst duration (Fig. 6h), average inter-burst interval (Fig. 6i), or average median inter-spike interval (ISI) within the burst (Fig. 6j). Collectively, these results indicate that *TSC2* variants lead to hyperactive neuronal network activity in TSC organoids, which may contribute to epilepsy in the TSC-affected brain.

### Aberrant synapse and neurofilament formation and abnormal mitochondria in TSC brain

To corroborate the pivotal observations in TSC cortical organoids, specifically the mitochondrial abnormalities and elevated neurofilament levels, we performed TEM imaging on surgically excised cortical tissue samples from both TSC-affected and normal brains. Similar to TSC organoids, we observed abnormal mitochondria in TSC brain, including enlarged and autophagic mitochondria (Fig. 7a). The size of the mitochondria significantly increased in the TSC brain (Fig. 7b). Neurofilaments observed in the TSC surgically excised cortical tissue samples demonstrated tight, intertwined bundles of filamentous aggregates that were running the length of the axon (Fig. 7c). The axonal plasma membranes enveloping the densest of these bundles often appeared loose, or ‘disconnected’ from the filamentous structures within the axon, causing the axonal membranes to take on an undulating, ‘wavy’ morphology (Fig. 7c). To confirm the increase of neurofilament in cortical tissue samples from TSC patients, we performed NEFL immunostaining (Fig. 7d). Quantitative analysis revealed a significantly increased percentage of total NEFL^+^ area relative to the entire image area in TSC compared to controls (Fig. 7e). In addition, we quantified the density of synaptic puncta in human brain tissues by immunocytochemistry of SYN1 and dendrite marker MAP2 (Fig. 7f). We found that the density of SYN1 synaptic puncta was significantly increased in the TSC brain (Fig. 7g), which is consistent with the observations in TSC organoids. Importantly, the qPCR analyses validated the increase in the NLGN and NRXN families in the TSC brain, including *NLGN1*, *NLGN3*, *NRXN1*, and *NRXN3* (Fig. 7h-k), which is consistent with our transcriptomic and proteomic results in TSC organoids. Consequently, our results not only corroborate the utility of human iPSC-derived brain organoids as an authentic model for examining TSC pathophysiology but also furnish compelling evidence that aberrations in synapses, neurofilaments, mitochondria, and NLGN1 are intricately linked with TSC.

**Figure 7.**
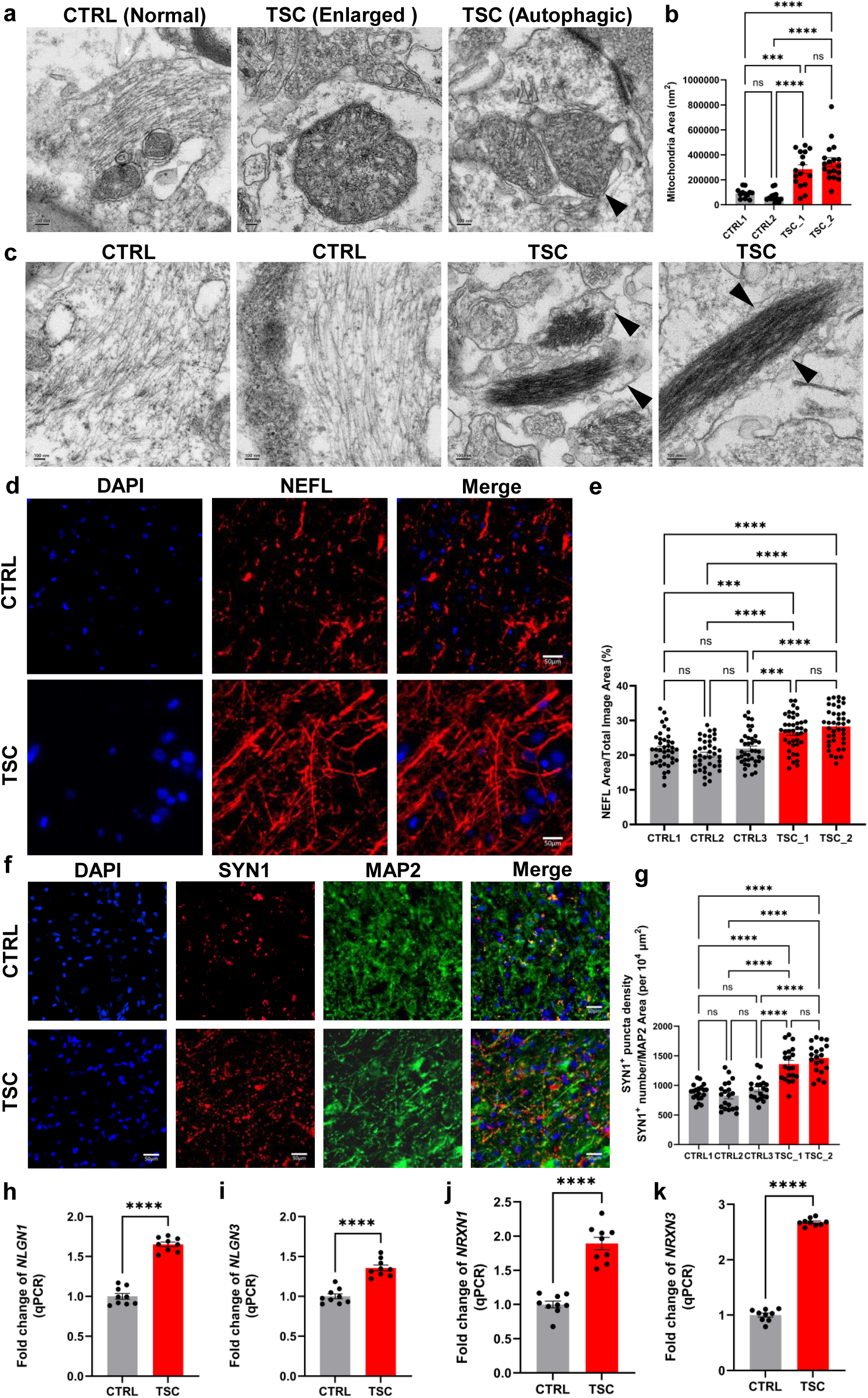
Aberrant synapse and neurofilament formation and abnormal mitochondria in TSC brain. **(a)** Ultrastructure of mitochondria from control and TSC surgically excised cortical tissues. Mitochondria residing in a neuron from control surgically excised cortical tissue (left); enlarged mitochondria in neurons from TSC surgically excised cortical tissue (middle); an arrowhead indicates the double-layered membrane of an autophagic vacuole engulfing two mitochondria in the TSC surgically excised cortical tissue. Scale bars, 100 nm. **(b)** Quantification of mitochondrial size in control and TSC surgically excised cortical tissue. Data are presented as mean ± s.e.m. (n = 12-13 mitochondria from each CTRL subject and n = 16-18 from each TSC patient were analyzed, \*\*\**P* < 0.001, \*\*\*\**P* < 0.0001, one-way ANOVA). **(c)** Ultrastructure of axons from control and TSC surgically excised cortical tissues. Ultrastructure of unmyelinated axons from control surgically excised cortical tissue (left); ultrastructure of axons from TSC surgically excised cortical tissue (right); black arrowheads denote the wavy/loose axonal membranes enveloping dense bundles of neurofilaments found in TSC surgically excised cortical tissues. Scale bars, 100 nm. **(d-e)** The percentage of NEFL^+^ area relative to the entire image area is significantly increased in TSC brain. Representative images (d) and stereological quantification (e) are presented, showing NEFL^+^ (red) in the total image area. Data are presented as mean ± SEM. A total of 40 views were analyzed from each control and TSC surgically excised cortical tissues. \*\*\**P* < 0.001, \*\*\*\**P* < 0.0001, one-way ANOVA), scale bars, 50 μm. **(f-g)** *TSC2* mutations accelerate synapse formation. Shown are sample images (f) and quantification of SYN1^+^ (g) puncta density in both control and TSC surgically excised cortical tissues. scale bars, 50 μm. Data are presented as mean ± s.e.m. (n = 20 views in each sample were analyzed from three control and two TSC surgically excised cortical tissues, \*\*\*\**P* < 0.0001, one-way ANOVA). **(h-k)** qPCR of *NLGN1*, *NLGN3*, *NRXN1* and *NRXN3* mRNA expression in the control and TSC human brain. Data are presented as mean ± s.e.m. (\*\*\*\**P* < 0.0001, unpaired t test).

## DISCUSSION

Although substantial advancements have been achieved in comprehending TSC signaling pathways in animal models, the precise effects of pathogenic *TSC1/2* variants on the progression of human brain development are still largely unclear. In this study, by conducting a thorough and longitudinal multi-omics and cellular analyses of TSC organoids, we demonstrate for the first time that pathogenic *TSC2* variants disrupt the neurodevelopmental trajectories through perturbations of gene regulatory networks and cell-cell communications during early cortical development, leading to mitochondrial abnormalities, aberrant neurofilament formation, and impaired synaptic formation and neuronal network activity. Importantly, the key findings in TSC cortical organoids, particularly the mitochondrial abnormalities, elevated neurofilament levels, and increased synapse formation were corroborated in surgically removed cortical tissues from TSC-affected brains. This validation underscores the significance of human iPSC-derived brain organoids as effective models for studying neuropsychiatric disorders.

Brain development is characterized by a series of dynamic, distinct phases. Through longitudinal transcriptomic investigations, we can examine the modulation of gene regulatory networks by *TSC2* variants across developmental stages. This insight is vital for delineating the specific temporal and consequential impacts of these genetic modifications. We identified the highest number of DEGs on day 35 (Fig. 1c-d), suggesting that the impact of *TSC2* variants on the disease originates at an early stage. This knowledge is imperative for the strategic timing of therapeutic interventions to coincide with the most vulnerable phases of cerebral development. Strikingly, the differentially expressed genes that keep increasing during development in TSC organoids are strongly associated with nervous system development and synaptic transmission. Indeed, TSC organoids exhibited accelerated differentiation of excitatory neurons, increased synapse formation, and hyperactive neuronal network activity. Conversely, the differentially expressed genes that were concomitantly decreasing in TSC organoids are largely enriched in cell cycle, DNA repair, and chromosomal functions (Fig. 1i-j). Therefore, *TSC2* variants significantly disrupt the developmental trajectories of the central nervous system. Mutations in the *TSC2* gene are implicated in the pathogenesis of TSC, predisposing individuals to a spectrum of neurological manifestations such as ASDs, epilepsy, intellectual disabilities, and TSC-associated neuropsychiatric disorders (TAND). Additionally, we identified that downregulated DEGs in TSC organoids are enriched in cytosolic ribosomes (Fig. 3f), while downregulated DEPs in synaptosomes of TSC are enriched in mitochondrial ribosomes (Fig. 5f). Furthermore, NLGN1 target genes include several ribosomal protein genes (Fig. 4e-f). Ribosomal impairments and deficits in ribosome production have been observed in a range of neurodegenerative diseases, including Alzheimer’s, Huntington’s, amyotrophic lateral sclerosis (ALS), and frontotemporal lobe dementia (FTLD) ^46–49^. These disruptions affect protein synthesis, contributing to the cellular dysfunction and degeneration characteristic of these disorders. This suggests that reduced ribosomal biogenesis may also contribute to the onset and progression of neurodevelopmental disorders. Our longitudinal transcriptomic analyses reveal marked disparities in the temporal expression patterns of genes linked to neurodevelopmental and neuropsychiatric disorders within TSC organoids. These findings illuminate the molecular pathways affected by such variants, providing a deeper understanding of the pathophysiology underlying tuberous sclerosis complex.

One hallmark we observed in TSC organoids is the increase of gliogenesis, in particular, reactive astrocytes. Indeed, analyses of postmortem tissues have revealed a pronounced accumulation of reactive astrocytes within the cortical and subcortical white matter regions of brains affected by TSC ^50^. Yet, the question remains whether the proliferation of reactive astrocytes in the TSC-afflicted brain arises as a secondary effect of cerebral lesions, including cortical tubers, subependymal nodules, and subependymal giant cell astrocytomas, or stems from mechanisms independent of tuber/tumor formation. Previous studies noted that astrocytes lacking *TSC2* expression or derived from TSC patients exhibit a higher saturation density and manifest enhanced proliferative behavior relative to control astrocytes ^12,13^. Aligned with these studies, we observed increased levels of NNAT and S100B, markers for early-stage and mature astrocytes, respectively, in TSC organoids throughout development. Furthermore, cell type composition analysis conducted on single-cell RNA sequencing datasets from TSC organoids corroborates the elevation of reactive astrocytes within these organoids (Fig. 3b). Considering the absence of cortical tubers/tumors in TSC organoids during these developmental stages, our results imply that the proliferation of reactive astrocytes in brains affected by TSC originates intrinsically from the gene regulatory networks driven by *TSC2* pathogenic variants, independent of tuber/tumor formation. This indicates a fundamental pathological process at play, distinct from the structural abnormalities traditionally associated with TSC, offering new insights into the disease’s etiology and potential therapeutic targets.

Astrocytes are integral to the overall homeostasis of the brain’s microenvironment, influencing neuronal development and health, synaptic communication, and, ultimately, cognitive functions ^51–53^. Neuron-astrocyte interactions are essential not only for the formation and maintenance of synapses but also for their dynamic modulation in response to neuronal activity ^54^. In this study, we demonstrated, for the first time, that *TSC2* pathogenic variants disrupt the crosstalk between neurons and reactive astrocytes through the neuroligin (NLGN)-neurexin (NRXN) signaling network. NLGN1, a high-confidence ASD risk gene ^55,56^, by interacting with neurexins or other related molecules, plays a pivotal role in synapse formation, stabilization, function, and maturation ^36,57–59^. The neuroligin signaling networks not only bridge pre- and post-synaptic neurons at synapses but also modulate astrocyte-neuron crosstalk which orchestrate functional synapse assembly ^36–40^. In mouse models, genetic modification of NLGNs results in ASD-like behaviors, along with a disrupted excitatory to inhibitory (E/I) balance ^60,61^. Notably, we identified a significant upregulation of *NLGN1* expression in TSC organoids and the TSC human brain, along with its binding partners including *NRXN1*, *NRXN3*, *CNTN4*, *CNTNAP2*, *CNTN1*, *CNTN2*, *CNTN3*, *CNTN5*, *CNTN6*, *CNTNAP4*, and *CNTNAP5* (Fig. 4a-b and Fig. 7h-k). In particular, the ligand-receptor pairs NLGN1-NRXN1, NLGN1-NRXN3, NLGN1-CNTNAP2, and NLGN1-CNTN4 are upregulated in TSC organoids. The ligand-receptor pairs NLGN1-NRXN1 and NLGN1-NRXN3 are increased in the TSC human brain. The upregulation of NLGN1 and other molecules in TSC organoids and TSC human brain may be mediated by the cap-dependent mRNA translation through the TSC1/2-mTORC1 pathway ^62^. Indeed, previous studies have demonstrated that enhanced cap-dependent translation results in an increase in mRNA translation and Nlgn1 protein expression in mice, correlating with ASD-related social behavior deficits ^62,63^. Through the cell-cell communication analyses, we further identified that the elevated NLGN1 signaling networks disrupt the neuron-astrocyte crosstalk in TSC organoids. Notably, the increased expression of *NLGN1* in TSC organoids could be rescued by rapamycin treatment. These observations underscore the significance of the NLGN1-mediated signaling network in the pathology of TSC, further positing the NLGN1-mediated signaling pathway as a viable therapeutic target for interventions in TSC.

Mitochondria are crucial in sustaining synaptic transmission, primarily through the generation of ATP and the regulation of intracellular calcium levels within the synapse. Dysfunctions or defects in mitochondrial operations are poised to exert broad impacts across various aspects of synaptic plasticity ^64,65^. This underscores the mitochondria’s critical role in supporting the dynamic processes underpinning synaptic adaptation and strength, highlighting how mitochondrial anomalies can disrupt the delicate balance of neural network function and plasticity. In this study, we found impaired mitochondrial translation integrity and mitophagy in synapses of TSC organoids and surgically excised cortical tissues from TSC-affected brains. Our results are consistent with previous studies showing impaired mitochondrial homeostasis and mitophagy in *Tsc1/2*-deficient neurons and mouse models ^18^. Indeed, mitochondrial dysfunction has been demonstrated to contribute to the etiology of neuropsychiatric disorders such as autism spectrum disorders ^66,67^, highlighting the importance of mitochondrial health in maintaining neural function.

Among the most striking and novel characteristics observed in both TSC organoids and brains is the pronounced increase in neurofilament formation. Within the neurons of the central nervous system (CNS), neurofilaments are structured as heteropolymers composed of four subunits: NEFL (neurofilament light chain), NEFM (neurofilament medium chain), NEFH (neurofilament heavy chain), and either α-internexin or peripherin. ^29–31^. Neurofilaments, as intermediate filaments, primarily provide structural support to neuronal axons, ensuring their integrity and facilitating efficient signal transmission ^30^. Neurofilament subunits are also located in postsynaptic terminal boutons, suggesting they may have a role in modulating neurotransmission, potentially influencing synaptic efficiency and plasticity ^30^. Among neurofilaments, NEFL has been identified as a biomarker in a range of neurological disorders ^68^. For example, a previous study observed increased NEFL levels in both cerebrospinal fluid (CSF) and blood in diseases of the CNS and peripheral nervous system linked to axonal injury or degeneration ^69^. Notably, the cohort with TSC displayed increased levels of NEFL in CSF in comparison to healthy control individuals ^70^. In this study, we provided the first direct evidence of elevated NEFL levels in both TSC organoids and brain specimens from TSC patients, highlighting a potential biomarker for TSC-related neural pathology. Future research is required to more precisely define the impact of elevated neurofilament levels on the pathogenesis of TSC, specifically regarding its involvement in TSC-associated neuropsychiatric disorders and epilepsy.

## ACKNOWLEDGEMENTS

We are deeply grateful to Dr. Gary J. Bassel from the Department of Cell Biology and Dr. Nisha Raj from the Department of Human Genetics at Emory University School of Medicine for their support and expert guidance. Their contributions were instrumental in the successful completion of this work. This work was supported in part by Department of Defense grant (W81XWH1910353 to Z.W. and M.J.G.), NIH grants (R01AG065611, R01MH121102, RF1AG079256, and R21MH132012 to Z.W.; RF1AG064909 to J.P.), and the Emory Integrated Genomics Core Facility (RRID:SCR_023529). This study was also supported by the Robert P. Apkarian Integrated Electron Microscopy Core (IEMC) at Emory University, which is subsidized by the School of Medicine and Emory College of Arts and Sciences. Additional support for IEMC is provided by the Georgia Clinical & Translational Science Alliance of the National Institutes of Health under award number UL1TR000454. Transmission electron micrographs were collected on the JEOL JEM-1400, 120kV TEM supported by the National Institutes of Health Grant S10 RR025679 or the Hitachi.

## AUTHOR CONTRIBUTIONS

W.N. led and was involved in all aspects of the project. W.N. performed all the cell culture experiments and multi-well multielectrode arrays and analyses. W.N. wrote the initial manuscript draft. S.Y. and W.N. performed transcriptomics and single cell transcriptomics analyses. W.N. and S.Y. contributed to preparing figures for transcriptomics, proteomics, and single cell transcriptomics. W.N. and X.L. contributed to immunocytochemistry, imaging, and their analysis. Z.W. performed proteomics and their preliminary analyses. R.C. helped with data analyses. W.N. and C.M. contributed to flow cytometry. W.N. and T.J.W contributed to TEM. X.L. helped with sample collection. A.J., Y.Z., and J.J.C. collected the surgically resected cortical specimens. M.J.G obtained IRB approval, consented the families and collected urine samples from TSC patients. J.P. designed proteomics. Z.Q., B.L., M.J.G. and Z.W. supervised this project. Z.W. revised the manuscript. Z.W. and W. N. designed the project and finalized the manuscript.

## Data Availability

RNA-seq and scRNA-seq data have been deposited in the Gene Expression Omnibus (NCBI) under accession number GSE247369. Similarly, the mass spectrometry proteomics data are available in the ProteomeXchange Consortium via the PRIDE partner repository, with the dataset identifier PXD046379. All these data will be made available at the time of publication.

## Code Availability

Code will be available at the time of publication.

## METHODS

### Subjects and iPSC generation

Urine samples (100-200 ml) from four TSC patients carrying *TSC2* variants (Supplementary Table 1) were collected at the Children’s Healthcare of Atlanta Tuberous Sclerosis Complex (TSC) Clinic. This was followed by the isolation of exfoliated renal epithelial cells. Induced pluripotent stem cells (iPSCs) were then generated using the CytoTune™-iPS 2.0 Sendai Reprogramming Kit (ThermoFisher) according to the manufacturer’s instructions and were fully characterized, including karyotyping, genotyping, expression of pluripotency markers, and in vitro differentiation capability. All studies adhered to institutional IRB protocols approved by the Emory University School of Medicine, and informed consent was obtained from all subjects or their legal representatives. A control iPSC line (PGP1) was edited to separately introduce homozygous TSC009 *TSC2* variant (c.C3643T), heterozygous and homozygous *TSC2* variant (c.G4514C) using CRISPR-Cas9 gene editing (Synthego) to generate three TSC isogeneic lines.

Research involving human surgically excised cortical tissue from 3 non-TSC control subjects and 2 TSC patients (Supplementary Table 9) was approved by the Emory University Institutional Review Board. Each TSC patient had been diagnosed with *TSC2* variants. The procedure was fully explained to the subjects or their legal representatives, all of whom provided signed informed consent.

### Human induced pluripotent stem cell-derived cortical organoids cultures

All iPSC lines were maintained on plates coated with Matrigel (Corning) and cultured in human iPSC mTeSR™1 media (Stemcell Technologies). Cortical organoids were differentiated from iPSCs using a protocol modified from previous publications ^19^. Briefly, intact human iPSC colonies were collected by Collagenase IV (Gibco) and moved to a 10 cm suspension culture dish, with daily changes of embryonic (EB) media. EB media contains DMEM/F12 (Corning), 20% KnockOut Serum Replacement (ThermoFisher), 1X GlutaMAX™ (ThermoFisher), 1X MEM non-essential amino acids solution (Invitrogen, 11140-050), 0.1 mM beta-mercaptoethanol (ThermoFisher), 2 μM A-83, and 2 μM Dorsomorphin dihydrochloride (Tocris/fisher). On day 6, media were replaced with neural media (NM) featuring Neurobasal (Invitrogen), 1X B-27 serum substitute without vitamin A (Invitrogen), 1X GlutaMAX™, 100 U/ml penicillin and streptomycin (Invitrogen). This NM was supplemented with 20 ng/ml bFGF (Peprotech) and 20 ng/ml EGF (peprotech), with daily media changes for the initial 10 days and alternate days for the following 9 days. On day 25, to promote differentiation of the neural progenitors into neurons, we replaced bFGF and EGF with 20 ng/ml BDNF (Peprotech). The media was changed every other day with NM boosted with BDNF. On day 43, media were replaced with growth factor-free neural medium, and the media was changed every four days. This medium included Neurobasal, 1X B-27 serum substitute without vitamin A, 1X GlutaMAX™, 100 U/ml penicillin and streptomycin. However, on day 70, the media - which was changed every four days - included 1X B-27 serum with vitamin A (Thermo).

### RNA isolation, qPCR and bulk RNA sequencing

Organoids were thoroughly homogenized using TRIzol Reagent (Invitrogen) and then followed as the manufacturer’s guidelines. Subsequently, RNA was extracted utilizing the Direct-zol RNA Miniprep Kit (Zymo). qPCR was performed using the Luna Universal One-Step RT-qPCR Kit (NEB, E3005E), following the manufacturer’s instructions. Each reaction was performed in triplicate and analyzed using the ΔΔCt method, with glyceraldehyde-3-phosphate dehydrogenase (GAPDH) serving as the normalization control. The primers were summarized in supplementary Table 10.

An RNA-seq library was constructed using the Illumina mRNA sample prep kit (cat. no. RS-100-0801) as directed by the manufacturer. To isolate poly-A-containing mRNA, poly-T oligo-attached magnetic beads were employed. This mRNA was subsequently fragmented into small segments using divalent cations at elevated temperatures. These RNA fragments were then transformed into first-strand cDNA using reverse transcriptase and random primers. The synthesis of the second-strand cDNA involved DNA Polymerase I and RNase H. These cDNA fragments underwent a terminal repair process where an ‘A’ base was added, and an adapter was ligated. Following gel purification, the final cDNA libraries were constructed and enriched through PCR. Prior to sequencing on the Illumina NovaSeq6000, the library constructs’ size and concentration were confirmed using a bioanalyzer. The quality of the raw reads was examined to ensure the efficacy of both library preparation and sequencing for further analysis. Adapter sequences, contaminant sequences (such as poly-A tails), and low-quality sequences with PHRED quality scores below 5 were trimmed from the reads using fastq tools as necessary ^71^. The RNA-seq reads were then aligned to the human reference genome hg38 using STAR v2.10. Differential expression values were calculated with DESeq2 v1.26.0. Genes characterized by an FDR < 0.05 and a log2FoldChange exceeding 1.5 or falling below -1.5 were identified as differentially expressed genes (DEGs). Enrichment analyses of different gene sets were performed with R package gprofiler2 (v0.2.2, https://cran.r-project.org/web/packages/gprofiler2/index.html) and the results were filtered by FDR < 0.05. Protein-protein interaction networks were extracted from the R package STRINGdb ^44^ with a confidence score > 400. R 4.2.0 was used for all downstream analyses, including plotting and statistical tests.

### Immunocytochemistry

Whole organoids or human brain tissues were fixed in 4% paraformaldehyde (PFA) (VWR) for 1-2 hours at room temperature. Samples were then washed with PBS for three times and submerged in a 30% sucrose solution overnight at 4℃. These samples were subsequently embedded in optimal cutting temperature (OCT, Fisher) and sectioned using a Leica cryostat. For the immunostaining, tissue samples were washed with PBS thrice. Samples were permeabilized with 0.2% Triton-X (Sigma) in PBS for 30 minutes and blocked with 10% donkey serum and 0.1% Triton-X in PBS for 30 minutes at room temperature. The following primary antibodies were used: anti-CTIP2 (rat, 1:500; Abcam, ab18465), anti-MAP2 (chicken, 1:500; Novus, nb300-213), anti-SYN1 (mouse, 1:500; Synaptic Systems, 106011), anti-PSD95 (rabbit, 1:500; Thermo Fisher, 51-6900), anti-NNAT (rabbit, 1:500; Abcam, ab27266), anti-S100B (rabbit, 1:500; Thermo Fisher, 710363) and anti-NEFL (rabbit, 1:500; Abcam, ab223343). Secondary antibodies used for immunostaining include donkey anti-mouse Alexa Fluor 488 (1:1000; ThermoFisher #R37114), donkey anti-goat IgG (H+L) Alexa Fluor 568 (1:1000; ThermoFisher #A-11057), donkey anti-rat Alexa Fluor 488 (1:1000; ThermoFisher #A21208), donkey anti-chicken Cy5 (1:1000; Jackson ImmunoResearch #703-175-155), donkey anti-rabbit Alexa Fluor 568 (1:1000; ThermoFisher #A10042), and Hoechst 33342 (1:2,000; Thermo Fisher, 62249). Antibodies were prepared in PBS containing 0.1% Triton X-100, 5% donkey serum (Sigma). Sections were mounted using Fluoromount-G Slide Mounting Medium (SouthernBiotech). Imaging was conducted with a Nikon Eclipse Ti-E microscope, and the ImageJ software from the National Institutes of Health was employed to tally cells showing positive results for the markers. The staining method for human surgically excised cortical tissue is the same as the above-mentioned method for organoid staining.

### FACS and Flow Cytometer Analysis

Organoids were dissociated into single-cell suspensions using the Papain Dissociation System (Worthington Biochemical). The cell viability exceeded 98%. The NNAT antibody (Abcam, ab27266) was conjugated with PE/R-Phycoerythrin using the Conjugation Kit (Abcam, ab102918) as per the manufacturer’s instructions. Following this, the cells were treated with the Foxp3/Transcription Factor Staining Buffer Set (Invitrogen, 00552300). The detailed procedures are outlined below: Prepare the fixation buffer and permeabilization buffer. Cells were fixed in 200 μL fixation buffer for 20 minutes at room temperature. Then cells were permeabilized with 1 mL of permeabilization buffer and centrifuged at 500g for 5 minutes at room temperature. Cells were resuspended in 200 μL of permeabilization buffer containing the previously prepared NNAT antibody for 1 hour. Add 1 mL of permeabilization buffer and centrifuge at 500g at room temperature for 5 minutes. Discard the supernatant. For flow cytometry analysis, cells were incubated with 2% PFA in PBS and acquired on a BD Symphony with at least 15,000 live cells obtained for each condition. Analysis was performed with FlowJo v10.8.1.

### MEA recording

CytoView MEA 6-well (Axion, M384-tMEA-6B) were coated with 0.1% polyethyleneimine (PEI) (Sigma-Aldrich, P3143) and 10 μg/ml laminin (Sigma, L2020). Organoids were seeded on MEA plates and maintained in organoid media at 37°C and 5% CO_2_. Two weeks after seeding, MEA recordings were performed. Spontaneous neuronal activities were recorded using a Maestro Edge MEA system and AxIS software (Axion BioSystems). The recordings were conducted with Neural Broadband. For spike detection, the threshold was set to six times the rolling standard deviation of the filtered field potential on each electrode. Ten-minute recordings were analyzed to calculate the average spike rate and the number of active electrodes per well. An active electrode was defined as spike rate ≥ 0.5/min.

### Single cell RNA sequencing (scRNA-seq)

Single cell RNA-seq was conducted in 3 TSC lines (TSC007, TSC010, and TSC010HO) and 3 control lines (C1-2, 426, and PGP1) on day 98. For each cell line, there are two batches, with one organoid per cell line in each batch of scRNA-seq data. The first batch includes 3 controls (C1-A: C1-2_A; C2-A: 426_A; C3-A: PGP1_A) and 3 TSC patients (T1-A: TSC007_A; T2-A: TSC010_A; T3-A: TSC010HO_A). The second batch also includes 3 controls (C1-B: C1-2_B; C2-B: 426_B; C3-B: PGP1_B) and 3 TSC patients (T1-B: TSC007_B; T2-B: TSC010_B; T3-B: TSC010HO_B). Organoids were dissociated into single-cell suspensions using Papain Dissociation System (Worthington Biochemical, LK003153) as directed by the manufacturer. Cell viability was higher than 98%.

The raw sequencing files were processed using CellRanger (v7.1.0) from 10X Genomics to obtain the expression matrix for each sample. The reference genome used for raw read alignment was GRCh38, also obtained from 10X Genomics. To perform single-cell analysis, Seurat (V4.0) ^72^ was utilized. The filtered expression matrix files, generated by the CellRanger pipeline, were imported as Seurat objects. Quality control steps were then performed on the data. Cells with a number of genes between 200 and 10,000 and with less than 20% mitochondrial gene expression were retained. Any doublet cells were identified and removed using the R package scDblFinder ^73^.

To integrate all the TSC and control samples, we created a list of Seurat objects and applied the following steps to each object: NormalizeData, FindVariableFeatures, ScaleData, and RunPCA. Next, we selected 2000 integration features using SelectIntegrationFeatures. The integration anchors were identified using the preprocessed objects by applying FindIntegrationAnchors and Reciprocal PCA. After integration, we observed a significant batch effect in one of the samples. As a result, we excluded this sample from further analysis and performed all subsequent analysis on the integrated object. To identify cell clusters, we conducted UMAP analysis using the top 50 principal components. This analysis led to the identification of 13 distinct cell clusters. We then used the FindMarkers function in Seurat to identify marker genes for each cell cluster. Marker genes meeting the criteria of an adjusted *P* value < 0.05 and an absolute average log2FoldChange > 0.25 were considered significant and were used for cell type identification. Cell types were assigned to the clusters following modifications based on scRNA-seq data from previous studies ^25,26,74–76^ as follows: immature astrocytes (*HSPB1*), glial progenitor cells (*SOX2*, *HES1*, *HSPB1*, *SLC1A3*), deep layer excitatory neurons (*MAP2*, *BCL11B*, *RBFOX1*, *SLC17A6*, *SYT1*, *RELN*, *GRIN2B*, *TBR1*), astrocytes (*SLC1A3*, *SOX9*, *S100B*), upper layer excitatory neurons (*STMN1*, *STMN2*, *TUBB3*, *CUX1*, *POU3F2*, *SATB2*), mature interneurons (*GAD2*, *GAD1*, *SLC32A1*), intermediate progenitor cells (*ASCL1*, *PAX6*), immature preoptic area (pOA) interneurons (*SOX4*, *PAX7*, *MEF2C*, *ASXL3*, *LHX1*), reactive astrocytes (*SLC1A3*, *FABP7*), immature medial ganglionic eminence (MGE) interneurons (*DCX*, *CDH12*, *SYT1*, *PCP4*), proliferating neural progenitor cells (*MKI67*, *PAX6*, *SOX2*, *RPL35*, *SNRPD2*, *CENPF*, *TOP2A*, *HMGB2*, *PCNA*), radial glial cells (*HLA-B*, *FTL*, *HMOX1*), and outer radial glia cells (*SOX2*, *PAX6*, *HES1*, *HOPX*, *NES*, *FAM107A*, *FABP7*).

### RNA velocity Analysis

To obtain splicing-specific counts for each sample, we used Velocyto ‘run10x’ (V0.17) ^77^ based on the output of CellRanger. These counts were then used for downstream RNA velocity analysis. For the RNA velocity analysis, we utilized the scVelo toolkit (V0.2.5) ^78^. As scVelo is developed in Python and compatible well with Scanpy, we loaded the loom files (containing spliced and unspliced counts generated by Velocyto) along with the PCA and UMAP representations derived from Seurat in the previous steps into AnnData objects. We split the AnnData object into two separate objects, one for the TSC group and one for the control group. To prepare the dataset for analysis, we applied filtering and normalization using the scVelo function ‘pp.filter_and_normalize’. Additionally, we computed moments using ‘pp.moments’ to capture the transcriptional dynamics. Next, we used ‘tl.velocity’ with the mode set to ‘stochastic’ to compute cell velocities, which provides information on the direction and magnitude of cellular transitions. Additionally, we employed ‘tl.velocity_graph’ to compute a velocity graph, which further characterizes the relationship between cells based on their velocity information.

### Cell-Cell communication

We utilized the R package CellChat (V1.6.1) ^41^ to infer cell-cell interactions. Following the official workflow, we loaded the normalized counts into CellChat and applied several preprocessing functions with standard parameters. First, we used the ‘identifyOverExpressedGenes’ function to identify genes that were overexpressed in specific cell types. Then, with the ‘identifyOverExpressedInteractions’ function, we identified the overexpressed interactions between ligands and receptors. For our analysis, we used the CellChatDB.human database. Next, we used the ‘computeCommunProb’ function to infer the cell-cell communication network and filtered out communication if there were fewer than 100 cells in certain cell clusters. Additionally, we employed the ‘computeCommunProbPathway’ function to infer cell-cell communication at the signaling pathway level. To calculate the aggregated cell-cell communication network, we used the ‘aggregateNet’ function.

To identify target genes associated with the ligands involved in cell-cell communication patterns revealed by CellChat, we performed NicheNet ^42^ analysis using the R package nichenetr (V2.0.4). Specifically, we used Upper layer excitatory neuron and Deep layer excitatory neuron cells as sender cells, and Reactive astrocyte cells as receiver cells. We utilized the RNA assays from the integrated Seurat object and the ligand-receptor/ligand-target databases provided by nichenetr to calculate the ligand-receptor interactions and ligand-target interactions. The control data served as the reference condition, while the TSC data represented the condition of interest. Finally, we visualized the results of the ligand-target activity using the circlize package (V 0.4.15) ^79^.

### Synaptosome proteomics

On day 98, organoids were harvested from 3 TSC lines (TSC007, TSC010, and TSC010HO) and 3 control lines (C1-2, 426, and PGP1). Each cell line was represented in triplicate. Synaptosome proteins were extracted using the Syn-PER Synaptic Protein Extraction Reagent (Thermo, 87793). Sample preparation for proteomics was performed using the single-pot, solid-phase-enhanced sample-preparation (SP3) method following established procedures with modifications ^80^. In summary, the synaptosome protein pellet was lysed in 50 µL of SDS lysis buffer [2% SDS, 50 mM HEPES (pH 8.5), 5 mM DTT, 1X EDTA-free protease inhibitor, 1X PhosSTOP phosphatase inhibitor (Sigma-Aldrich)] and subjected it to sonication at 30% power for 30 seconds. The protein concentration in each sample was determined using a Coomassie-stained short SDS gel, as previously outlined ^81,82^. 25 µg of protein from each sample was used for proteomic analysis. First, the protein sample was reduced by fresh dithiothreitol (DTT, 1 mM) for 30 min, alkylated by iodoacetamide (IAA, 10 mM) at dark for 30 min, and further quenched with DTT (30 mM) for 30 min. Then 75 µg of SP3 beads was added to each sample, followed by adding absolute ethanol to achieve 50%. Protein binding was performed by incubating the sample in a ThermoMixer at 24 °C for 10 min at 1,000 r.p.m. After the binding was completed, the supernatant was removed, and the protein was washed three times using 200 µL of 80% ethanol. Finally, the protein was digested with Trypsin (Promega, 1:25 w/w) at 37°C overnight. The digested peptides of each sample were collected and dried using a Speedvac.

Each digested peptide sample was reconstituted in 50 mM HEPES (pH 8.5) to a concentration of 1 µg/µL, then labeled with 50 µg of TMTpro reagent (18-plex) for 30 minutes. Distinct TMT tags were used for individual samples. The labeling reaction was quenched using 0.5% hydroxylamine for 15 minutes. An equal amount of labeled samples were pooled and desalted with a Sep-Pak C18 Cartridge (Waters). 200 μg of desalted peptides underwent fractionation using offline basic pH RPLC with XBridge C18 columns (3.5 μm particle size, 2.1 mm × 15 cm, Waters). The process employed Buffer A (10 mM ammonium formate, pH 8.0) and Buffer B (90% AcN, 10 mM ammonium formate, pH 8.0). A 120-minute gradient of 15-50% buffer B was used to elute the peptides. Fractions were gathered every 0.5 minutes, subsequently concatenated into 40 fractions as previously outlined ^83^. Each concatenated fraction was dried and reconstituted in 5% formic acid for acidic pH LC-MS/MS analysis based on a refined protocol ^82^. In the LC-MS/MS analysis, a C18 column (75 μm × 20 cm, 1.9 µm C18 resin) was used with a 60 min gradient time (buffer A, 0.2% formic acid, 5% DMSO; buffer B, buffer A plus 65% acetonitrile) The settings of Q Exactive HF Orbitrap MS (Thermo Fisher Scientific). were as follows: MS1 scans: 60,000 resolution, 460-1600 m/z scan range, 1 x 10^6^ AGC, and 50 ms maximum ion time. 20 data-dependent MS2 scans: 60,000 resolutions, starting at 120 m/z, 1 x 10^5^ AGC, 120 ms maximum ion time, 1.0 m/z isolation window with 0.2 m/z offsets, HCD mode, 32% normalized collision energy, and a 10-second dynamic exclusion.

Protein identification and quantification were conducted using the JUMP search engine ^84^. The raw data were searched against the human protein database contains the downloaded Swiss-Prot, TrEMBL, and UCSC databases without redundancies(83,955 entries). A target-decoy approach was used to estimate the false discovery rate (FDR) ^85^. The primary search parameters included: A 15 ppm mass tolerance for both precursor and product ions, full tryptic restriction with up to two missed cleavages allowed, static modifications for TMT tags (+304.20715 on Lys and N-termini) and carbamidomethylation (+57.02146 on Cys), dynamic modifications for oxidation (+15.99491 on Met).. Post-search, peptide-spectrum matches (PSMs) were filtered based on mass accuracy and JUMP matching scores to achieve an FDR below 1%. In cases where peptides matched multiple homologous proteins, we attributed them to the protein with the most PSMs, adhering to the parsimony principle. Finally, proteins were quantified from the TMT reporter ion intensities, as previously detailed ^86^.

### Differentially expressed proteins and pathway enrichment analysis

To identify differentially expressed proteins (DEPs), we employed a methodology largely aligned with previous descriptions ^87^. Initially, we assessed experimental variances from replicated measurements, modeling them with a Gaussian distribution to determine the standard deviation (SD). The *P*-values for intergroup comparisons were derived from a moderated t-test, with FDR values calculated using the Benjamini-Hochberg (BH) correction to account for multiple testing. Criteria for DEPs were set at a change magnitude > 2 SD and an FDR < 0.05. We primarily probed our datasets for protein enrichments associated with synaptic compartments and processes using the SYNGO knowledgebase ^43^. Predominant protein modules were performed through the ClueGO (v2.5.10) gene ontology toolkit ^45^. Functional enrichment analyses of different protein sets were performed with R package gprofiler2 (v0.2.2, https://cran.r-project.org/web/packages/gprofiler2/index.html) and the results were filtered by FDR < 0.05. The PPI network was then visualized by R package STRINGdb ^44^ with confidence score > 400. R 4.2.0 was used for all downstream analyses, including plotting and statistical tests.

### Electron Microscopy

Organoids or human tissues were fixed in a mixture of 1.0% paraformaldehyde and 2.5% glutaraldehyde in 0.1M cacodylate buffer at pH 7.4 for overnight. The samples were post-fixed in 1.0% osmium tetroxide and subsequently stained en bloc with 2% aqueous uranyl acetate before being dehydrated in a stepwise manner through a series of graded ethanol concentrations up to 100% ethanol. Ethanol dehydration was followed by two changes in 100% propylene oxide. Samples were then embedded in Eponate 12 resin and polymerized at 60°C overnight. Ultrathin sections were cut with a Leica EM UC6 ultramicrotome, stained with uranyl acetate and lead citrate, and imaged in a Hitachi HT7700 TEM or a Jeol JEM 1400 TEM, both operated at 80 kV.

