## Supplementary Information for "Longitudinal multi-omics reveals pathogenic *TSC2* variants disrupt developmental trajectories of human cortical organoids derived from Tuberous Sclerosis Complex"

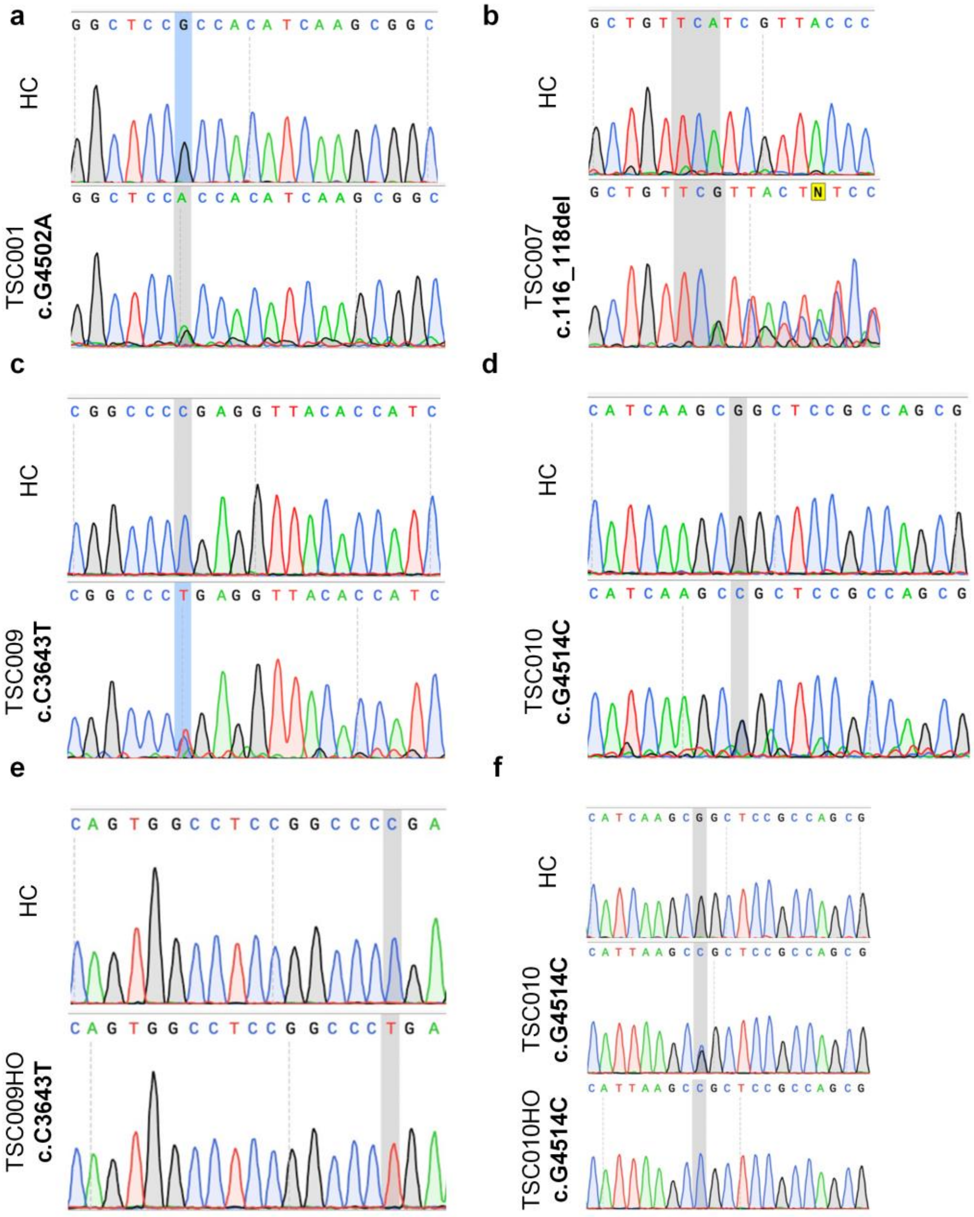


**Extended Data Figure 1. Genotyping TSC iPSC lines by Sanger sequencing.** The *TSC2* variants were confirmed through Sanger sequencing. The upper panels display the healthy control (HC) sequences, the lower panels highlight the *TSC2* variant sites. **(a)** TSC001: c.G4502A. **(b)** TSC007: c.116_118del. **(c)** TSC009: c.C3643T. **(d)** TSC010: c.G4514C. **(e)** TSC009HO: c.C3643T. **(f)** TSC010Het: c.G4514C and TSC010HO: c.G4514C. Het: Heterozygous. HO: Homozygous.

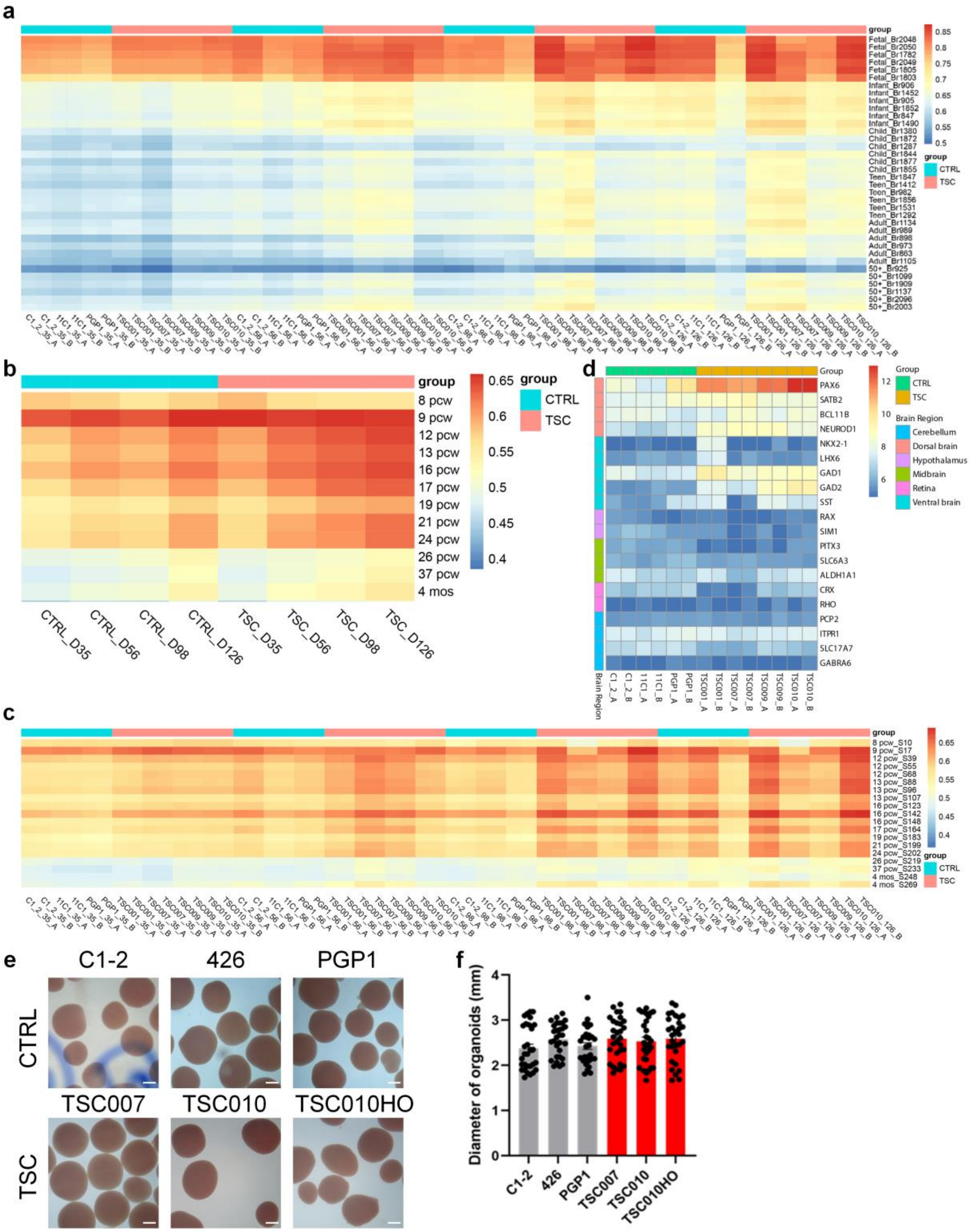
a
b
c
d
e
f
group
CTRL
TSC
Group
Brain Region
Cerebellum
Dorsal brain
Hypothalamus
Midbrain
Retina
Ventral brain
CTRL_D35
CTRL_D56
CTRL_D98
CTRL_D126
TSC_D35
TSC_D56
TSC_D98
TSC_D126
C1-2
426
PGP1
TSC007
TSC010
TSC010HO
Diameter of organoids (mm)

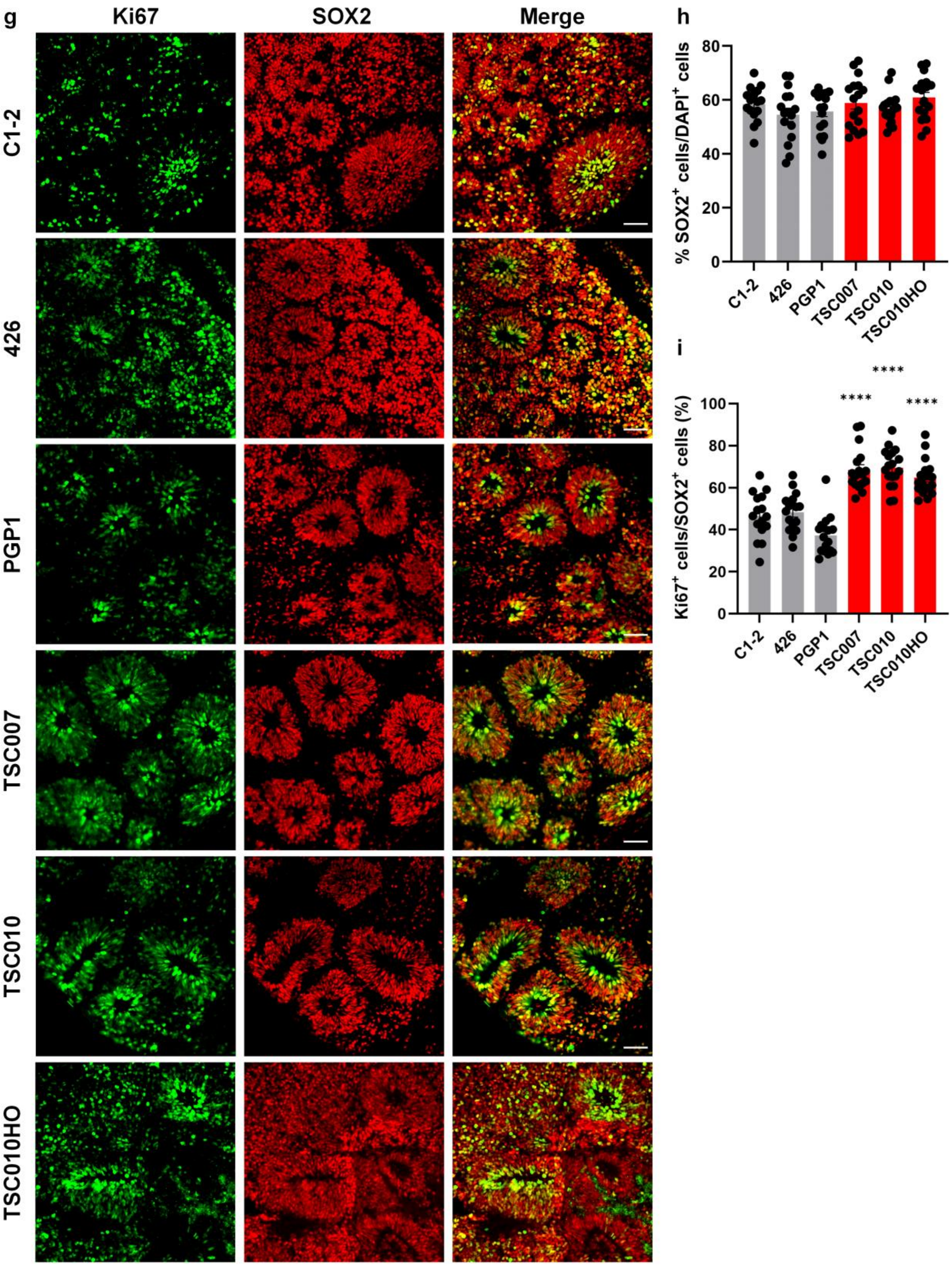
g
Ki67
SOX2
Merge
C1-2
426
PGP1
TSC007
TSC010
TSC010HO
h
% SOX2+ cells/DAPI+ cells
0
20
40
60
80
C1-2
426
PGP1
TSC007
TSC010
TSC010HO
i
Ki67+ cells/SOX2+ cells (%)
0
20
40
60
80
100
****
****
****
C1-2
426
PGP1
TSC007
TSC010
TSC010HO

**Extended Data Figure 2. Basic characterizations of TSC cortical organoids. (a)** Heatmap of Pearson's correlation analysis of RNA-seq datasets among cortical organoids at different stages and human dorsolateral prefrontal cortex (DLPFC) across 6 life stages (Jaffe et al., 2015). **(b)** Heatmaps of Pearson's correlation analysis of RNA-seq datasets among cortical organoids at different stages and published transcriptome datasets of human DLPFC at 11 developmental stages from BrainSpan. Colors indicate the averages for biological replicates. **(c)** Heatmap shows each sample's repeat of (b). **(d)** Heatmap shows the expression of marker genes specific to different brain regions in control and TSC cortical organoids. **(e)** Brightfield images of control and TSC organoids at day 98. Scale bar, 1 mm. **(f)** Quantification of organoid size in control and TSC groups. Data are presented as mean ± s.e.m. (n = 30 organoids per line, from three independent experiments; one-way ANOVA). **(g-i)** *TSC2* variants lead to increased NPC proliferation. Representative images (g) and quantifications illustrate the proportion of $SOX2^+$ NPCs among total $DAPI^+$ cells (h) and the proportion of $Ki67^+$ proliferating NPCs among $SOX2^+$ NPCs (i) in both control and TSC cortical organoids at Day 28. Data are presented as mean ± s.e.m. (n = 16-18 per cell line from three independent experiments). ****$P < 0.0001$, one-way ANOVA. Scale bars, 50 μm.

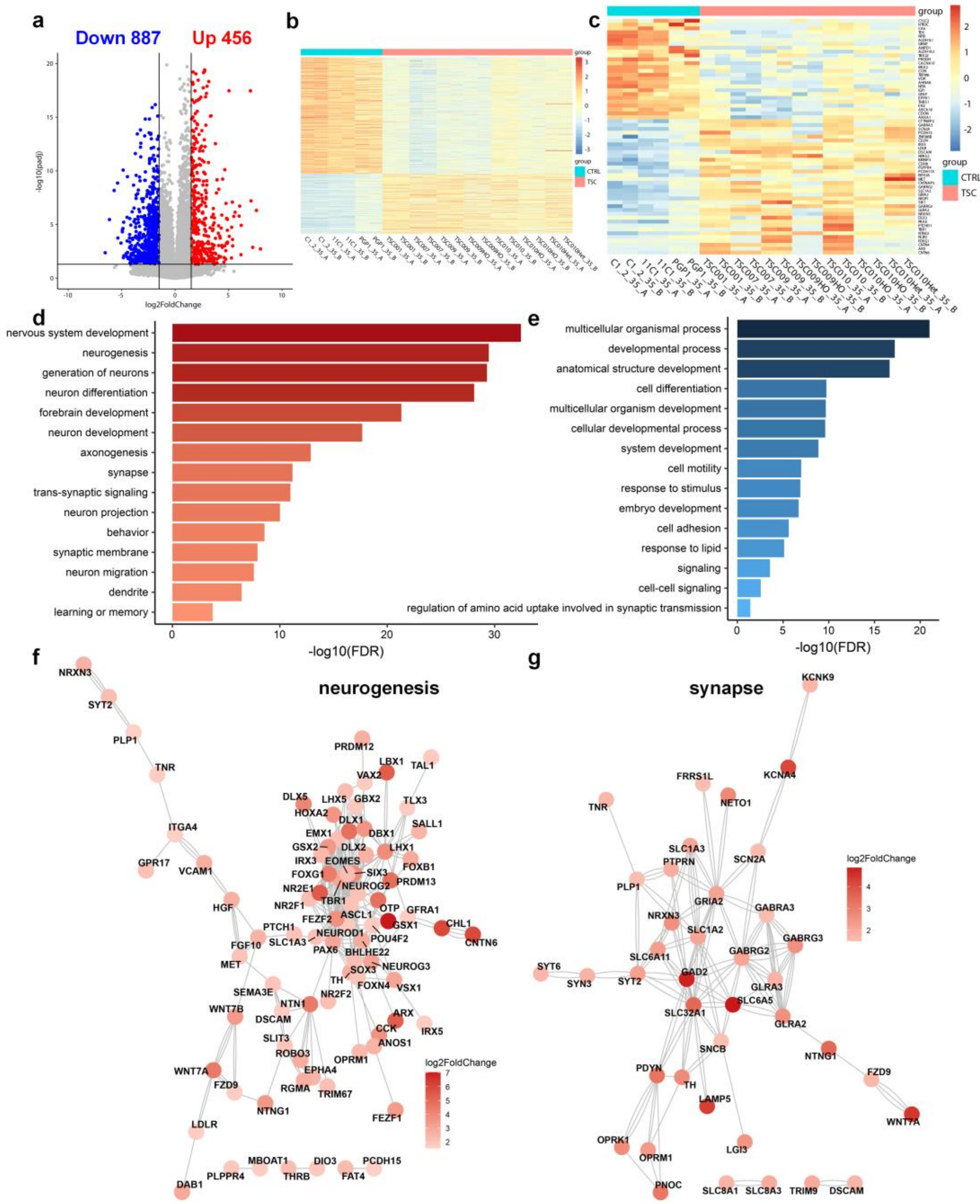


**Extended Data Figure 3. Transcriptomic analyses of TSC organoids at day 35. (a)** Volcano plot displays differentially expressed genes (DEGs) in TSC organoids at day 35 resulting from *TSC2* variants. Red and blue dots indicate 456 upregulated genes and 887 downregulated genes, respectively (adjusted *P*-value < 0.05, absolute log2 fold-change > 1.5). Grey dots represent genes with no significant differential expression. **(b)** Heatmap generated from DEGs in (a). **(c)** Heatmap of 61 overlapping genes between TSC DEGs and autism susceptibility genes from the SFARI Gene database. **(d-e)** Pathway analyses of upregulated (d) and downregulated (e) DEGs in TSC organoids. **(f-g)** Gene interaction analyses of the neurogenesis (f) and synapse (g) pathway reveal highly interactive functional network of DEGs in TSC organoids.

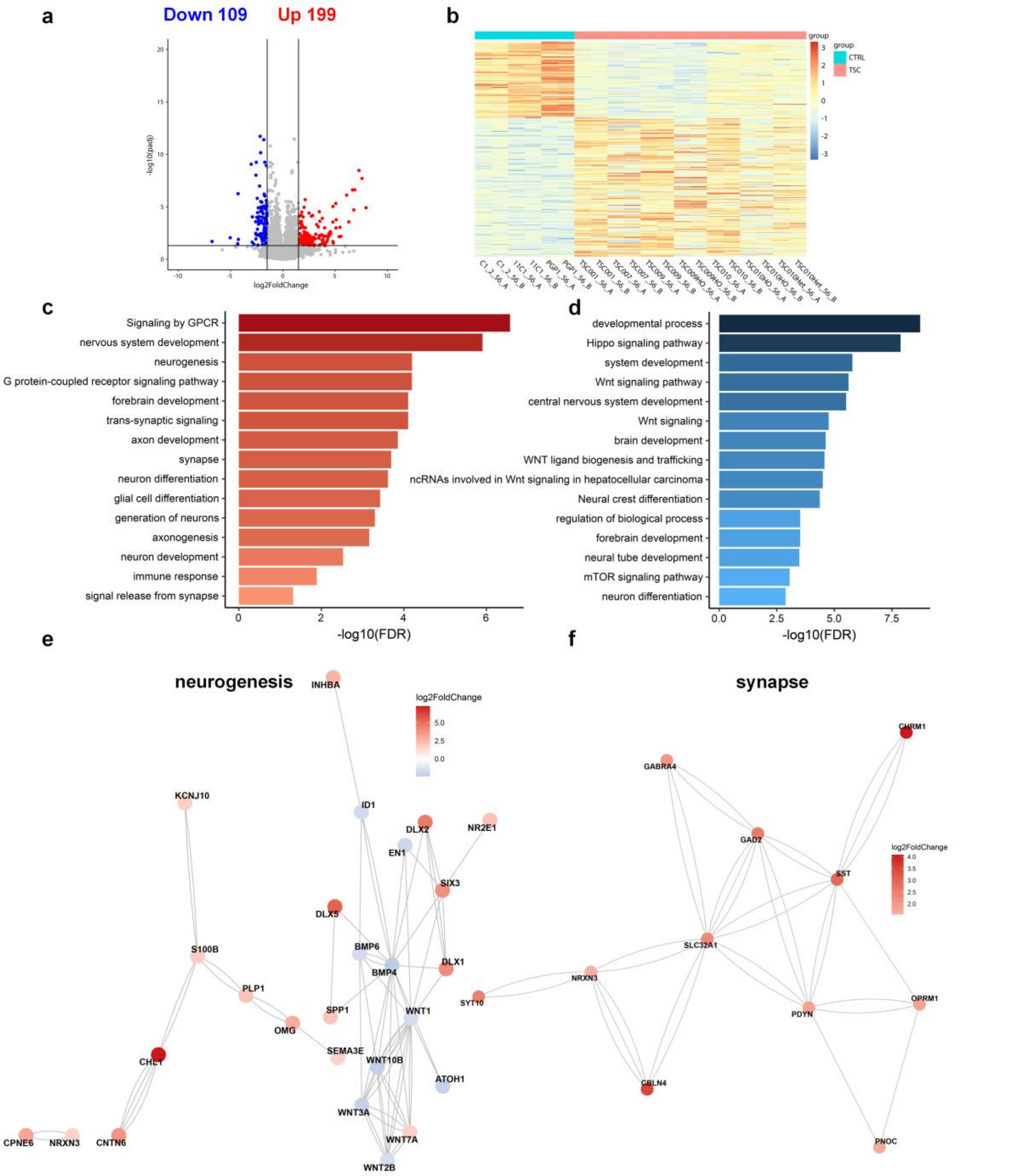


**Extended Data Figure 4. Transcriptomic analyses of TSC organoids at day 56. (a)** Volcano plot displays differentially expressed genes (DEGs) in TSC organoids at day 56 resulting from *TSC2* variants. Red and blue dots indicate 199 upregulated genes and 109 downregulated genes, respectively (adjusted *P*-value < 0.05, absolute log2 fold-change > 1.5). Grey dots represent genes with no significant differential expression. **(b)** Heatmap generated from DEGs in (a). **(c-d)** Pathway analyses of upregulated (c) and downregulated (d) DEGs in TSC organoids. **(e-f)** Gene interaction analyses of the neurogenesis (e) and synapse (f) pathway reveal highly interactive functional network of DEGs in TSC organoids.

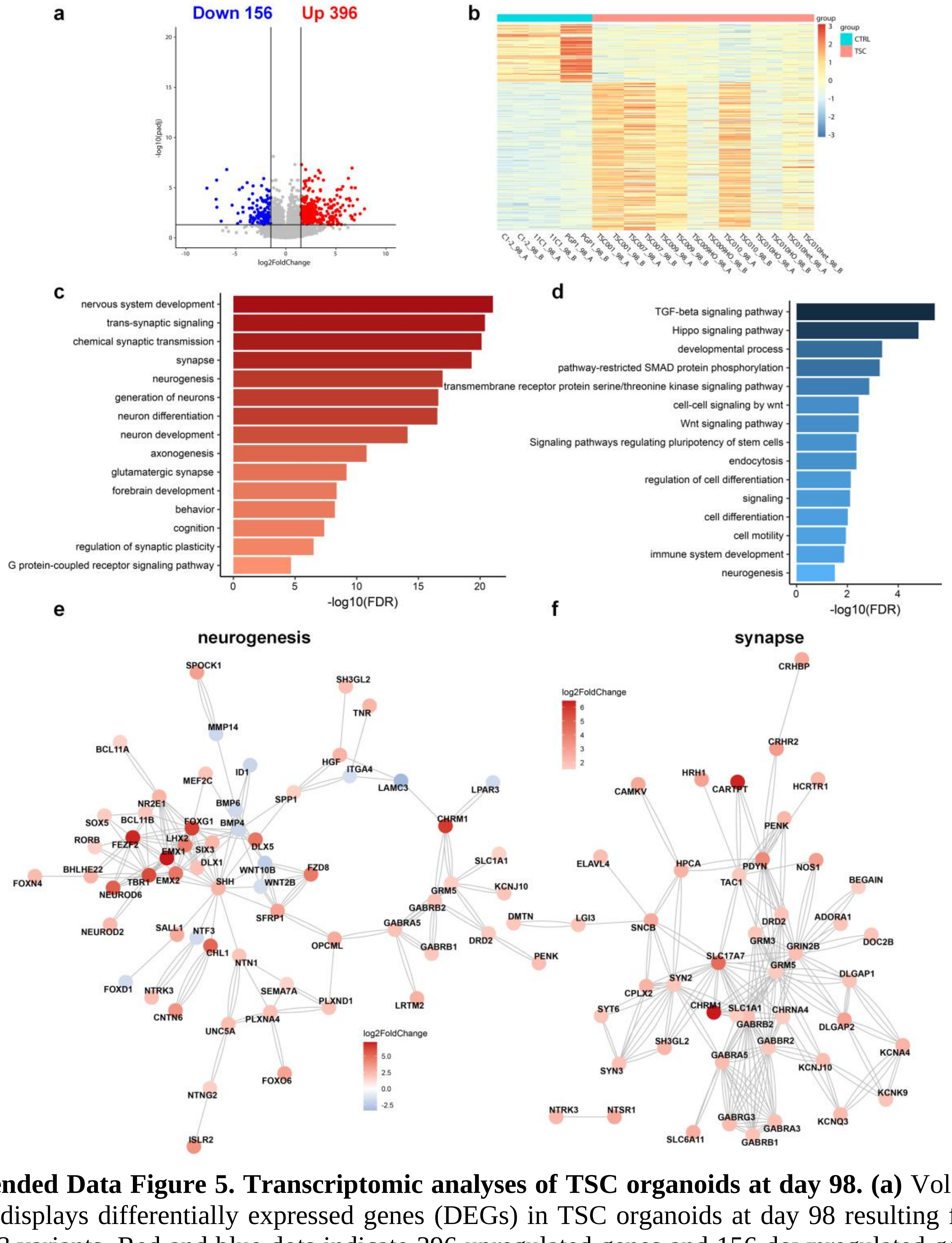


**Extended Data Figure 5. Transcriptomic analyses of TSC organoids at day 98. (a)** Volcano plot displays differentially expressed genes (DEGs) in TSC organoids at day 98 resulting from *TSC2* variants. Red and blue dots indicate 396 upregulated genes and 156 downregulated genes, respectively (adjusted *P*-value < 0.05, absolute log2 fold-change > 1.5). Grey dots represent genes with no significant differential expression. **(b)** Heatmap generated from DEGs in (a). **(c-d)** Pathway analyses of upregulated (c) and downregulated (d) DEGs in TSC organoids. **(e-f)** Gene interaction analyses of the neurogenesis (e) and synapse (f) pathway reveal highly interactive functional network of DEGs in TSC organoids.

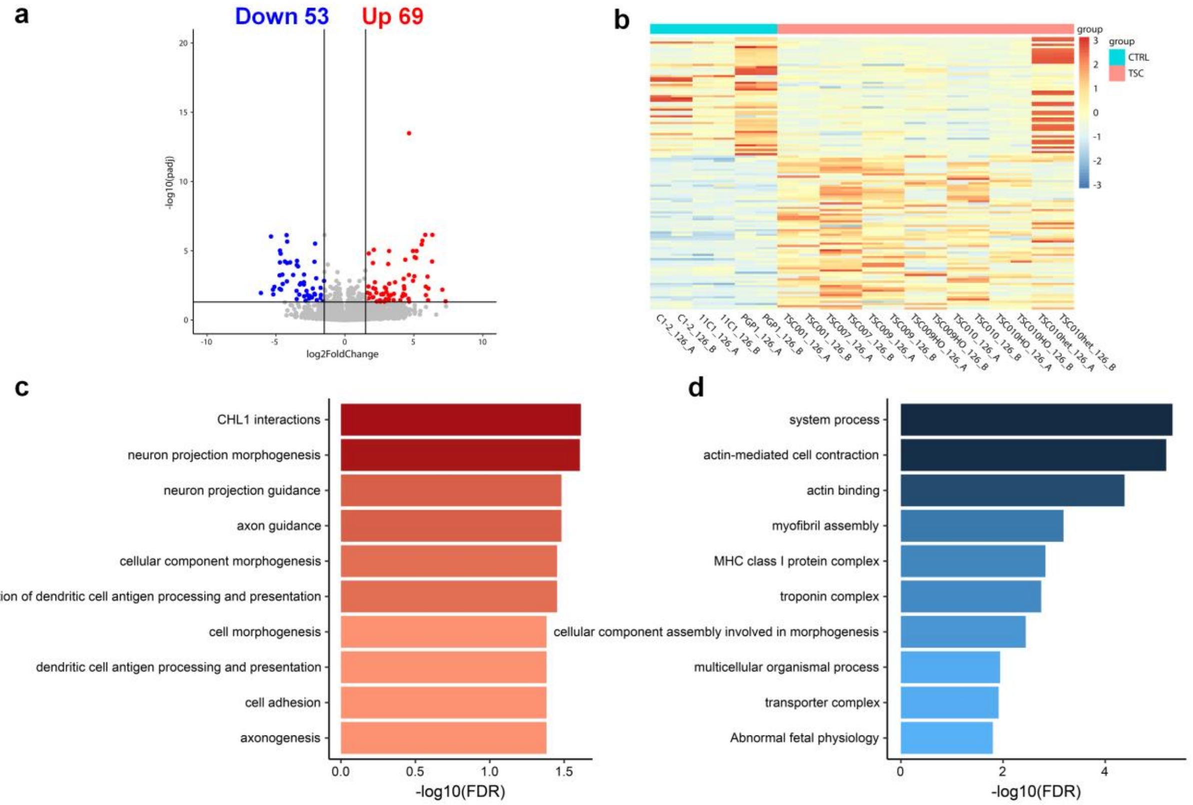


**Extended Data Figure 6. Transcriptomic analyses of TSC organoids at day 126. (a)** Volcano plot displays differentially expressed genes (DEGs) in TSC organoids at day 126 resulting from *TSC2* variants. Red and blue dots indicate 69 upregulated genes and 53 downregulated genes, respectively (adjusted *P*-value < 0.05, absolute log2 fold-change > 1.5). Grey dots represent genes with no significant differential expression. **(b)** Heatmap generated from DEGs in (a). **(c-d)** Pathway analyses of upregulated (c) and downregulated (d) DEGs in TSC organoids.

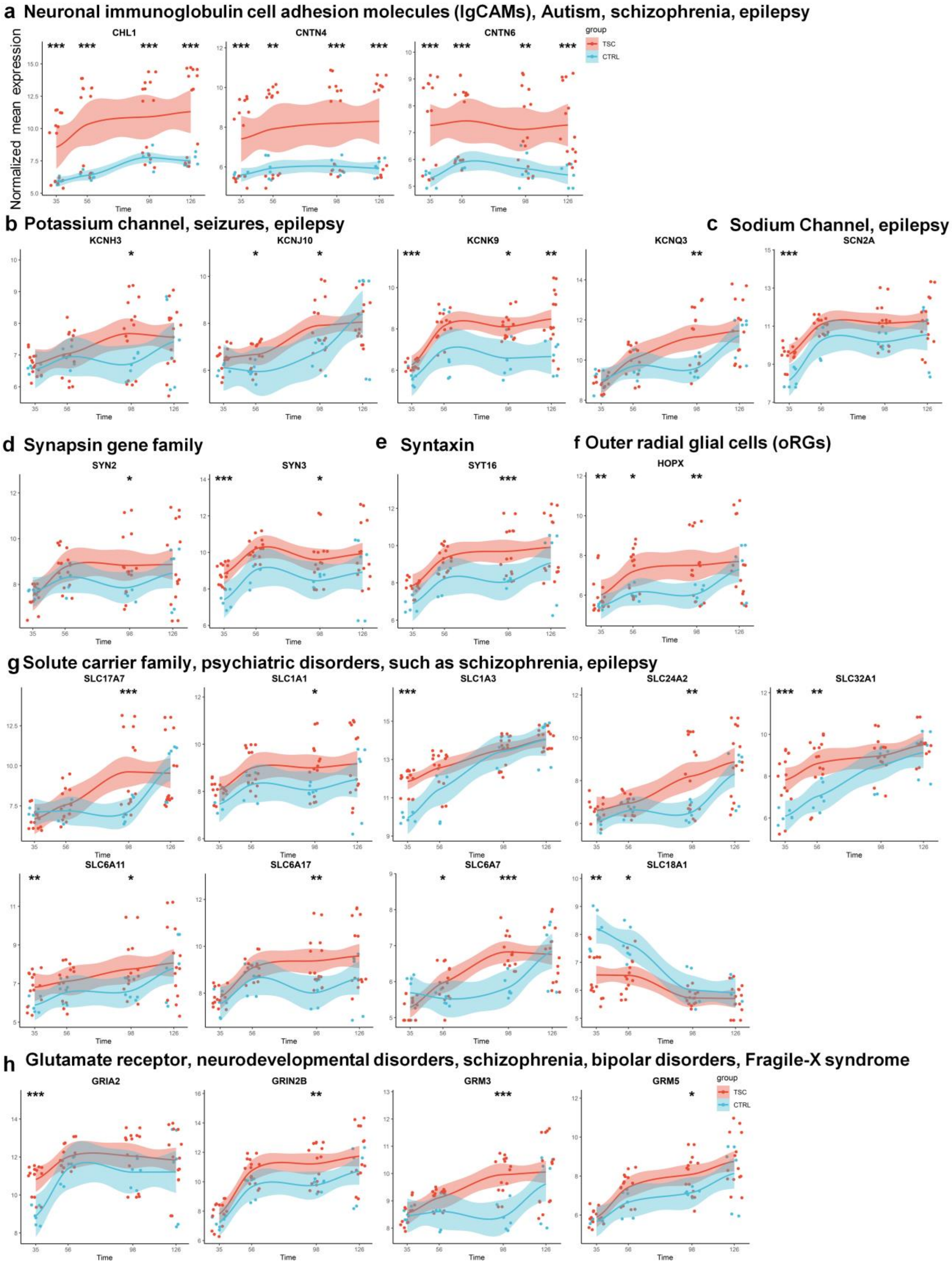
a Neuronal immunoglobulin cell adhesion molecules (IgCAMs), Autism, schizophrenia, epilepsy
CHL1
CNTN4
CNTN6
group
TSC
CTRL
Normalized mean expression
Time
b Potassium channel, seizures, epilepsy
KCNH3
KCNJ10
KCNK9
KCNQ3
c Sodium Channel, epilepsy
SCN2A
d Synapsin gene family
SYN2
SYN3
e Syntaxin
SYT16
f Outer radial glial cells (oRGs)
HOPX
g Solute carrier family, psychiatric disorders, such as schizophrenia, epilepsy
SLC17A7
SLC1A1
SLC1A3
SLC24A2
SLC32A1
SLC6A11
SLC6A17
SLC6A7
SLC18A1
h Glutamate receptor, neurodevelopmental disorders, schizophrenia, bipolar disorders, Fragile-X syndrome
GRIA2
GRIN2B
GRM3
GRM5
group
TSC
CTRL

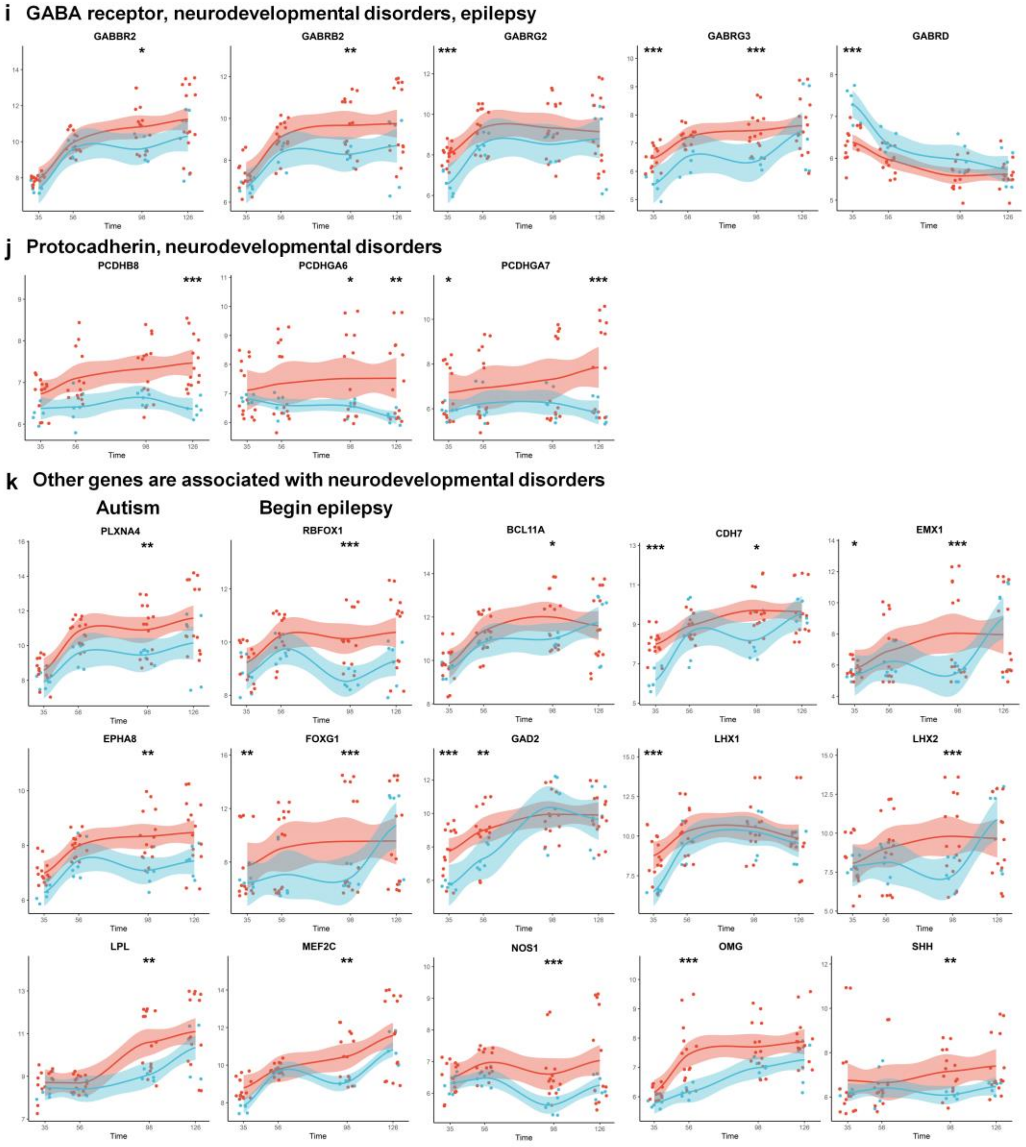


**Extended Data Figure 7. Altered expression trajectories of genes associated with neurodevelopmental disorders in TSC organoids.** Normalized mean expression of genes in the developmental trajectories in control and TSC organoids includes neuronal immunoglobulin cell adhesion molecules (IgCAMs) genes *CHL1*, *CNTN4*, and *CNTN6* **(a)**; potassium channel genes *KCNH3*, *KCNJ10*, *KCNK9*, and *KCNQ3* **(b)**; sodium channel gene *SCN2A* **(c)**; synapsin gene family *SYN2* and *SYN3* **(d)**; syntaxin gene *SYT16* **(e)**; Outer radial glial cells (oRGs) **(f)**; solute carrier family genes *SLC17A7*, *SLC1A1*, *SLC1A3*, *SLC24A2*, *SLC32A1*, *SLC6A11*, *SLC6A17*, *SLC6A7*, and *SLC18A1* **(g)**; glutamate receptor genes *GRIA2*, *GRIN2B*, *GRM3*, and *GRM5* **(h)**; GABA receptor genes *GABBR2*, *GABRB2*, *GABRG2*, *GABRG3*, and *GABRD* **(i)**; protocadherin genes *PCDHB8*, *PCDHGA6*, and *PCDHGA7* **(j)**; other genes associated with neurodevelopmental disorders *PLXA4*, *RBFOX1*, *BCL11A*, *CDH7*, *EMX1*, *EPHA8*, *FOXG1*, *GAD2*, *LHX1*, *LHX2*, *LPL*, *MEF2C*, *NOS1*, *OMG*, and *SHH* **(k)**. The red and blue area surrounding the trajectory line signifies the 95% confidence interval. * Adjusted $P$-value < 0.05, absolute log2 fold-change > 1.5; **$P$ < 0.01, absolute log2 fold-change > 1.5; ***$P$ < 0.001, absolute log2 fold-change > 1.5.

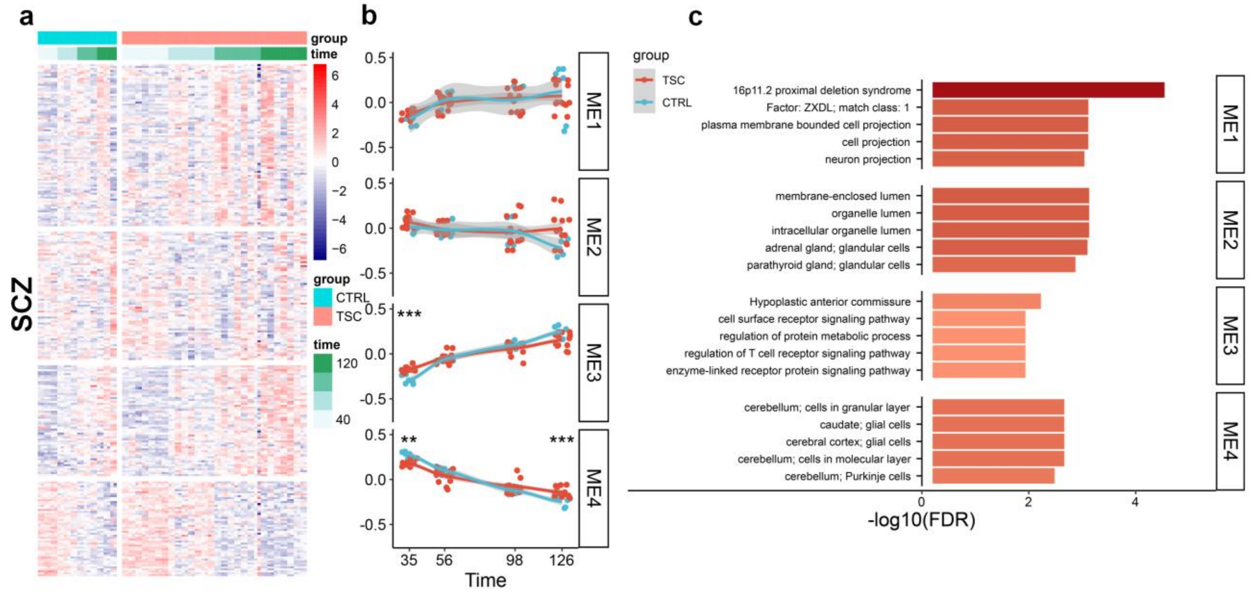


**Extended Data Figure 8. Mapping of schizophrenia genes onto the developmental trajectories of control and TSC organoid differentiation.** The initial column displays the grouping of standardized normalized expressions of genes linked to schizophrenia **(**SCZ; **a)**. The genes, arranged in rows, are grouped through hierarchical clustering based on the Euclidean distance between them. The samples, organized in columns, are sorted by the day of differentiation (indicated by green bars), starting with the earliest 35 days on the left and progressing to the latest time points 126 days on the right. The middle column **(b)** displays the first principal component of the eigengenes for the identified gene clusters. The grey area surrounding the trajectory line signifies the 95% confidence interval. $*P < 0.05$, $**P < 0.01$, $***P < 0.001$, two-sided Mann-Whitney test. The last column **(c)** presents the top GO terms that are enriched in the identified clusters.

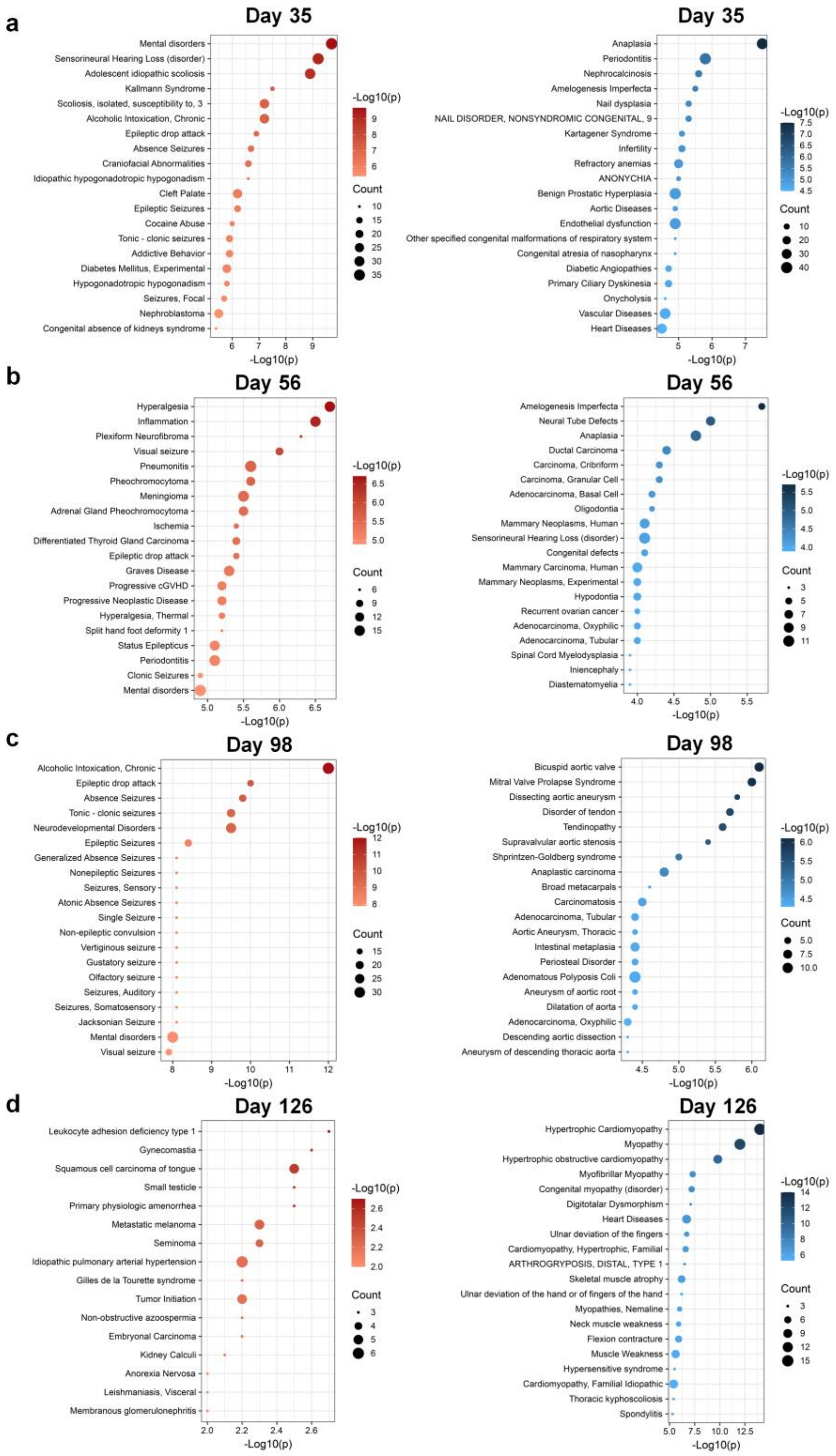


**Extended Data Figure 9. DisGeNET analyses DEGs of TSC organoids at four different stages.** Enrichment analyses were performed using DisGeNET for upregulated genes (red bubbles) and downregulated genes (blue bubbles) at day 35 **(a)**, day 56 **(b)**, day 98 **(c)**, and day 126 **(d)** (adjusted *P*-value < 0.05, absolute log2 fold-change > 1.5).

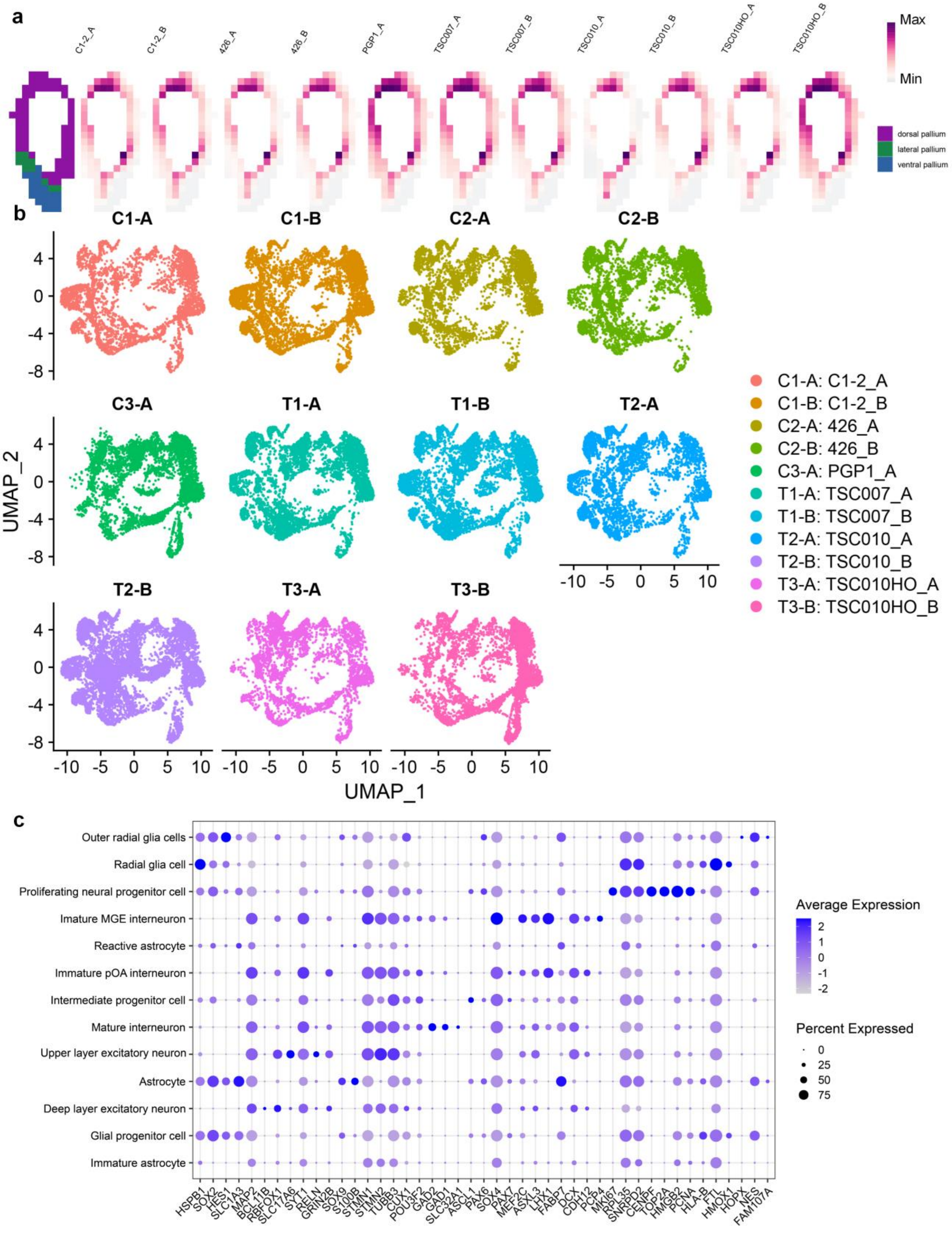
a
C1-2_A
C1-2_B
426_A
426_B
PGP1_A
TSC007_A
TSC007_B
TSC010_A
TSC010_B
TSC010HO_A
TSC010HO_B
Max
Min
dorsal pallium
lateral pallium
ventral pallium
b
C1-A
C1-B
C2-A
C2-B
C3-A
T1-A
T1-B
T2-A
T2-B
T3-A
T3-B
UMAP_2
UMAP_1
C1-A: C1-2_A
C1-B: C1-2_B
C2-A: 426_A
C2-B: 426_B
C3-A: PGP1_A
T1-A: TSC007_A
T1-B: TSC007_B
T2-A: TSC010_A
T2-B: TSC010_B
T3-A: TSC010HO_A
T3-B: TSC010HO_B
c
Outer radial glia cells
Radial glia cell
Proliferating neural progenitor cell
Imature MGE interneuron
Reactive astrocyte
Immature pOA interneuron
Intermediate progenitor cell
Mature interneuron
Upper layer excitatory neuron
Astrocyte
Deep layer excitatory neuron
Glial progenitor cell
Immature astrocyte
HSPB1 SOX2 HES1 SLC1A3 MAP2 BCL11B RBFOX1 SLC17A6 SYT1 RELN GRIN2B SOX9 S100B STMN1 STMN2 TUBB3 CUX1 POU3F2 GAD2 GAD1 SLC32A1 ASCL1 PAX6 SOX4 PAX7 MEF2C ASXL3 LHX1 FABP7 DCX CDH12 PCP4 MKI67 RPL35 SNRPD2 CENPF TOP2A HMGB2 PCNA HLA-B FTL HMOX1 HOPX NES FAM107A
Average Expression
Percent Expressed

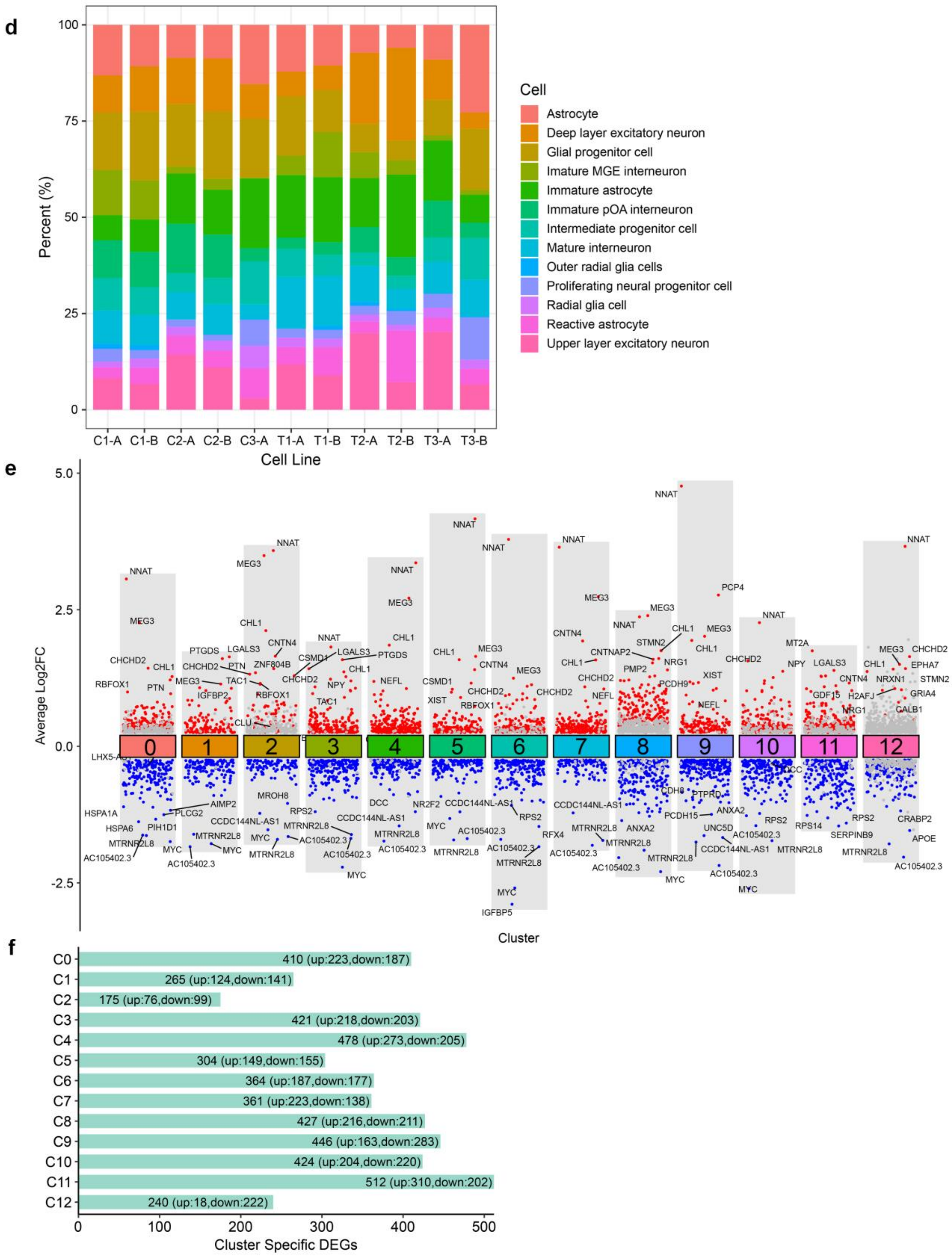

d
Percent (%)
100
75
50
25
0
C1-A
C1-B
C2-A
C2-B
C3-A
T1-A
T1-B
T2-A
T2-B
T3-A
T3-B
Cell Line
Cell
Astrocyte
Deep layer excitatory neuron
Glial progenitor cell
Imature MGE interneuron
Immature astrocyte
Immature pOA interneuron
Intermediate progenitor cell
Mature interneuron
Outer radial glia cells
Proliferating neural progenitor cell
Radial glia cell
Reactive astrocyte
Upper layer excitatory neuron
e
Average Log2FC
5.0
2.5
0.0
-2.5
0 1 2 3 4 5 6 7 8 9 10 11 12
Cluster
f
C0 410 (up:223,down:187)
C1 265 (up:124,down:141)
C2 175 (up:76,down:99)
C3 421 (up:218,down:203)
C4 478 (up:273,down:205)
C5 304 (up:149,down:155)
C6 364 (up:187,down:177)
C7 361 (up:223,down:138)
C8 427 (up:216,down:211)
C9 446 (up:163,down:283)
C10 424 (up:204,down:220)
C11 512 (up:310,down:202)
C12 240 (up:18,down:222)
0 100 200 300 400 500
Cluster Specific DEGs

**Extended Data Figure 10. Single cell transcriptomic analyses of TSC cortical organoids at day 98. (a)** VoxHunt plots show the expression similarity of cortical organoids derived from different iPSC lines to voxels in the developing mouse brain (embryonic day 13.5), along with the structural annotation of the sections. **(b)** UMAP projection of 98-day-old cortical organoids derived from each iPSC line used in this study. **(c)** Spot plot showing the expression of annotation reference genes across 13 cell clusters, as identified by scRNA-seq from day 98 control and TSC organoids. The spot color scale indicates the average expression for each gene, the spot size represents the percentage of cells in each cluster expressing the corresponding gene. **(d)** Bar plot showing the proportion of each cell type among all cells within each sample. **(e)** Volcano plot displays DEGs in TSC organoids compared to the controls. Red and blue dots represent upregulated and downregulated genes for each cell cluster, respectively (adjusted $P$-value < 0.05, absolute average log2 fold-change > 0.25). Grey dots represent genes with no significant differential expression. **(f)** DEG number of each cell cluster. Among the 13 clusters, radial glia cells (cluster 11) exhibited the highest number of DEGs.

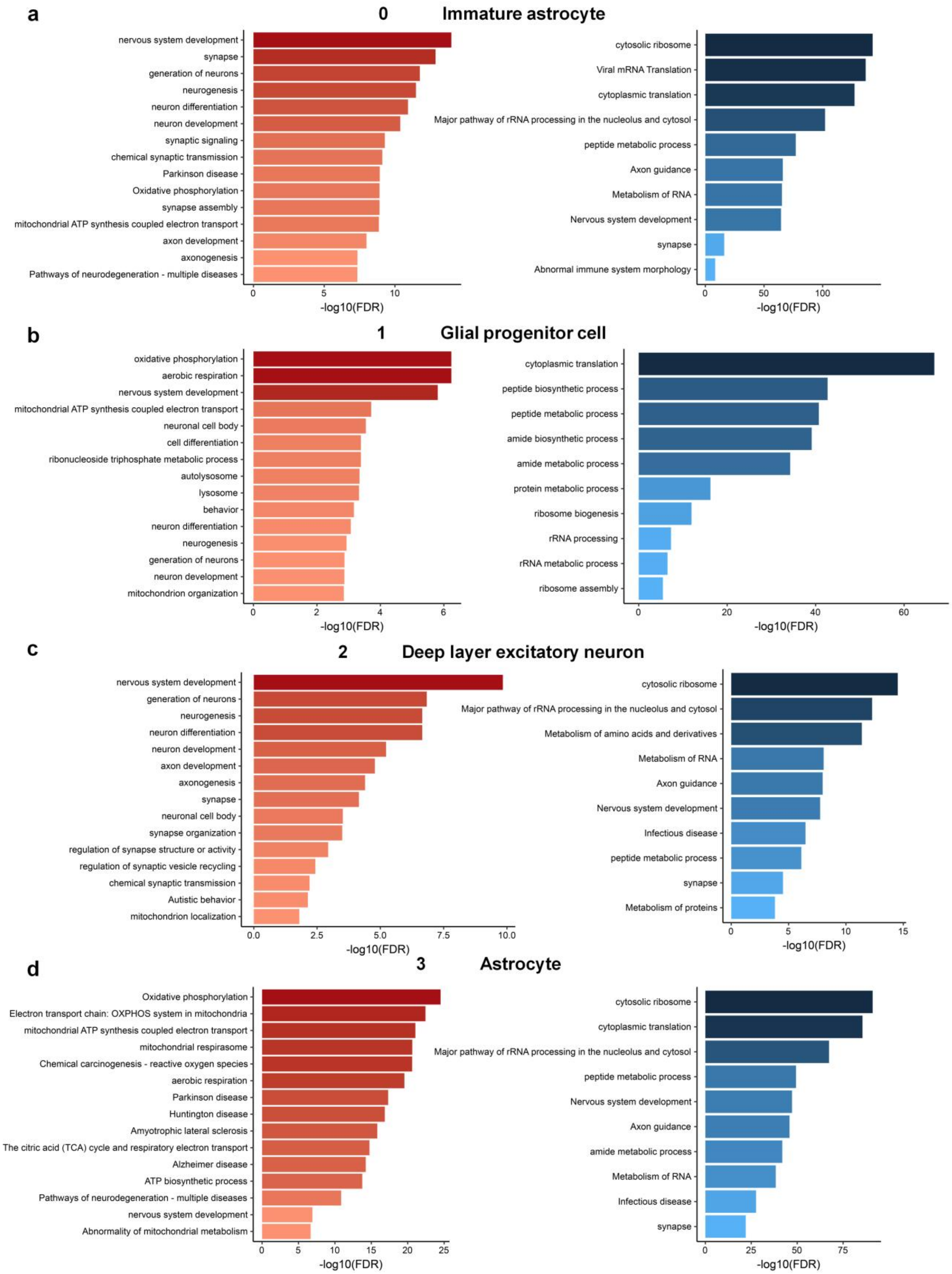
a
0 Immature astrocyte
nervous system development
synapse
generation of neurons
neurogenesis
neuron differentiation
neuron development
synaptic signaling
chemical synaptic transmission
Parkinson disease
Oxidative phosphorylation
synapse assembly
mitochondrial ATP synthesis coupled electron transport
axon development
axonogenesis
Pathways of neurodegeneration - multiple diseases
-log10(FDR)
cytosolic ribosome
Viral mRNA Translation
cytoplasmic translation
Major pathway of rRNA processing in the nucleolus and cytosol
peptide metabolic process
Axon guidance
Metabolism of RNA
Nervous system development
synapse
Abnormal immune system morphology
-log10(FDR)
b
1 Glial progenitor cell
oxidative phosphorylation
aerobic respiration
nervous system development
mitochondrial ATP synthesis coupled electron transport
neuronal cell body
cell differentiation
ribonucleoside triphosphate metabolic process
autolysosome
lysosome
behavior
neuron differentiation
neurogenesis
generation of neurons
neuron development
mitochondrion organization
-log10(FDR)
cytoplasmic translation
peptide biosynthetic process
peptide metabolic process
amide biosynthetic process
amide metabolic process
protein metabolic process
ribosome biogenesis
rRNA processing
rRNA metabolic process
ribosome assembly
-log10(FDR)
c
2 Deep layer excitatory neuron
nervous system development
generation of neurons
neurogenesis
neuron differentiation
neuron development
axon development
axonogenesis
synapse
neuronal cell body
synapse organization
regulation of synapse structure or activity
regulation of synaptic vesicle recycling
chemical synaptic transmission
Autistic behavior
mitochondrion localization
-log10(FDR)
cytosolic ribosome
Major pathway of rRNA processing in the nucleolus and cytosol
Metabolism of amino acids and derivatives
Metabolism of RNA
Axon guidance
Nervous system development
Infectious disease
peptide metabolic process
synapse
Metabolism of proteins
-log10(FDR)
d
3 Astrocyte
Oxidative phosphorylation
Electron transport chain: OXPHOS system in mitochondria
mitochondrial ATP synthesis coupled electron transport
mitochondrial respirasome
Chemical carcinogenesis - reactive oxygen species
aerobic respiration
Parkinson disease
Huntington disease
Amyotrophic lateral sclerosis
The citric acid (TCA) cycle and respiratory electron transport
Alzheimer disease
ATP biosynthetic process
Pathways of neurodegeneration - multiple diseases
nervous system development
Abnormality of mitochondrial metabolism
-log10(FDR)
cytosolic ribosome
cytoplasmic translation
Major pathway of rRNA processing in the nucleolus and cytosol
peptide metabolic process
Nervous system development
Axon guidance
amide metabolic process
Metabolism of RNA
Infectious disease
synapse
-log10(FDR)

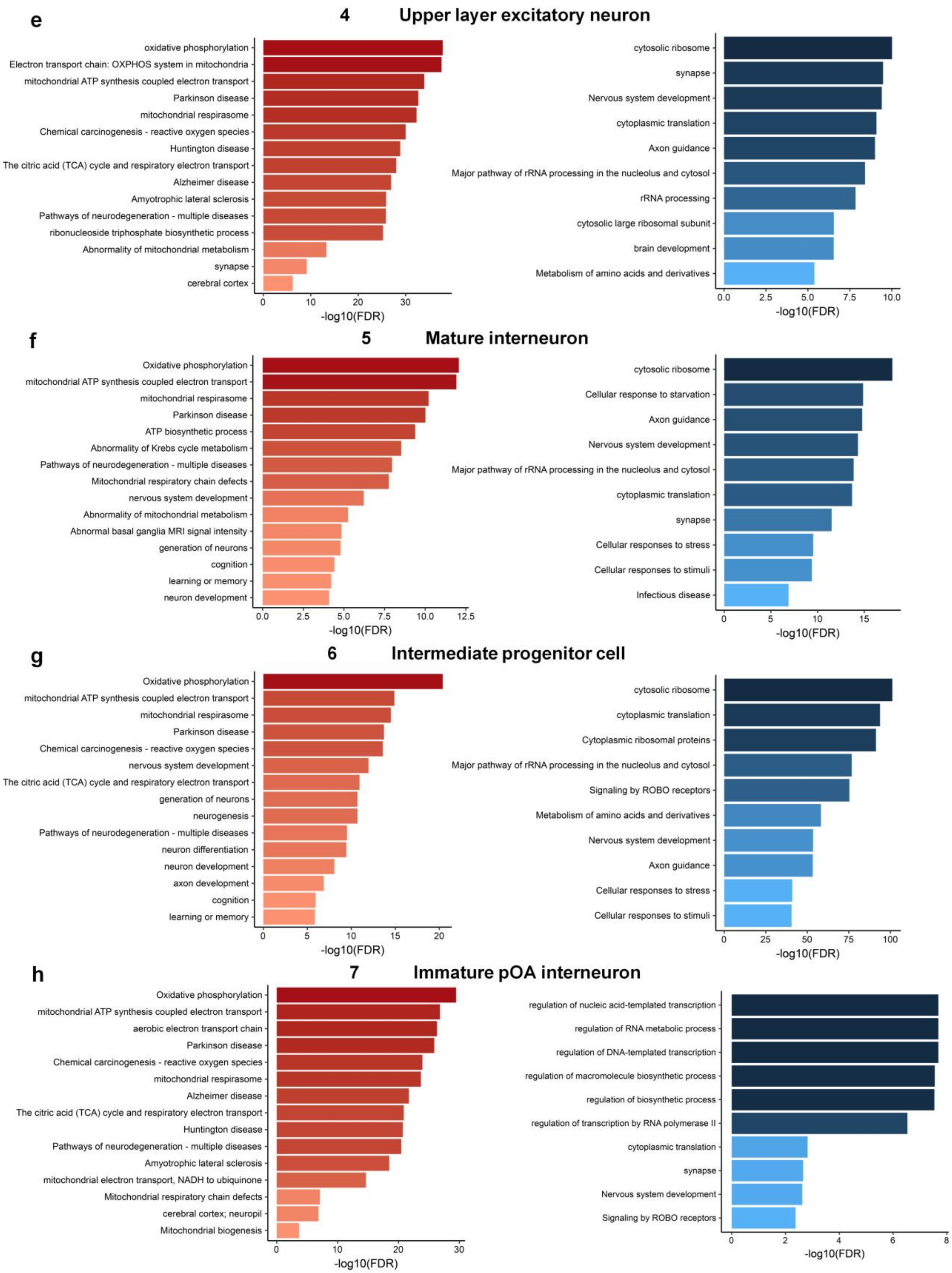
e
4 Upper layer excitatory neuron
oxidative phosphorylation
Electron transport chain: OXPHOS system in mitochondria
mitochondrial ATP synthesis coupled electron transport
Parkinson disease
mitochondrial respirasome
Chemical carcinogenesis - reactive oxygen species
Huntington disease
The citric acid (TCA) cycle and respiratory electron transport
Alzheimer disease
Amyotrophic lateral sclerosis
Pathways of neurodegeneration - multiple diseases
ribonucleoside triphosphate biosynthetic process
Abnormality of mitochondrial metabolism
synapse
cerebral cortex
-log10(FDR)
cytosolic ribosome
synapse
Nervous system development
cytoplasmic translation
Axon guidance
Major pathway of rRNA processing in the nucleolus and cytosol
rRNA processing
cytosolic large ribosomal subunit
brain development
Metabolism of amino acids and derivatives
-log10(FDR)
f
5 Mature interneuron
Oxidative phosphorylation
mitochondrial ATP synthesis coupled electron transport
mitochondrial respirasome
Parkinson disease
ATP biosynthetic process
Abnormality of Krebs cycle metabolism
Pathways of neurodegeneration - multiple diseases
Mitochondrial respiratory chain defects
nervous system development
Abnormality of mitochondrial metabolism
Abnormal basal ganglia MRI signal intensity
generation of neurons
cognition
learning or memory
neuron development
-log10(FDR)
cytosolic ribosome
Cellular response to starvation
Axon guidance
Nervous system development
Major pathway of rRNA processing in the nucleolus and cytosol
cytoplasmic translation
synapse
Cellular responses to stress
Cellular responses to stimuli
Infectious disease
-log10(FDR)
g
6 Intermediate progenitor cell
Oxidative phosphorylation
mitochondrial ATP synthesis coupled electron transport
mitochondrial respirasome
Parkinson disease
Chemical carcinogenesis - reactive oxygen species
nervous system development
The citric acid (TCA) cycle and respiratory electron transport
generation of neurons
neurogenesis
Pathways of neurodegeneration - multiple diseases
neuron differentiation
neuron development
axon development
cognition
learning or memory
-log10(FDR)
cytosolic ribosome
cytoplasmic translation
Cytoplasmic ribosomal proteins
Major pathway of rRNA processing in the nucleolus and cytosol
Signaling by ROBO receptors
Metabolism of amino acids and derivatives
Nervous system development
Axon guidance
Cellular responses to stress
Cellular responses to stimuli
-log10(FDR)
h
7 Immature pOA interneuron
Oxidative phosphorylation
mitochondrial ATP synthesis coupled electron transport
aerobic electron transport chain
Parkinson disease
Chemical carcinogenesis - reactive oxygen species
mitochondrial respirasome
Alzheimer disease
The citric acid (TCA) cycle and respiratory electron transport
Huntington disease
Pathways of neurodegeneration - multiple diseases
Amyotrophic lateral sclerosis
mitochondrial electron transport, NADH to ubiquinone
Mitochondrial respiratory chain defects
cerebral cortex; neuropil
Mitochondrial biogenesis
-log10(FDR)
regulation of nucleic acid-templated transcription
regulation of RNA metabolic process
regulation of DNA-templated transcription
regulation of macromolecule biosynthetic process
regulation of biosynthetic process
regulation of transcription by RNA polymerase II
cytoplasmic translation
synapse
Nervous system development
Signaling by ROBO receptors
-log10(FDR)

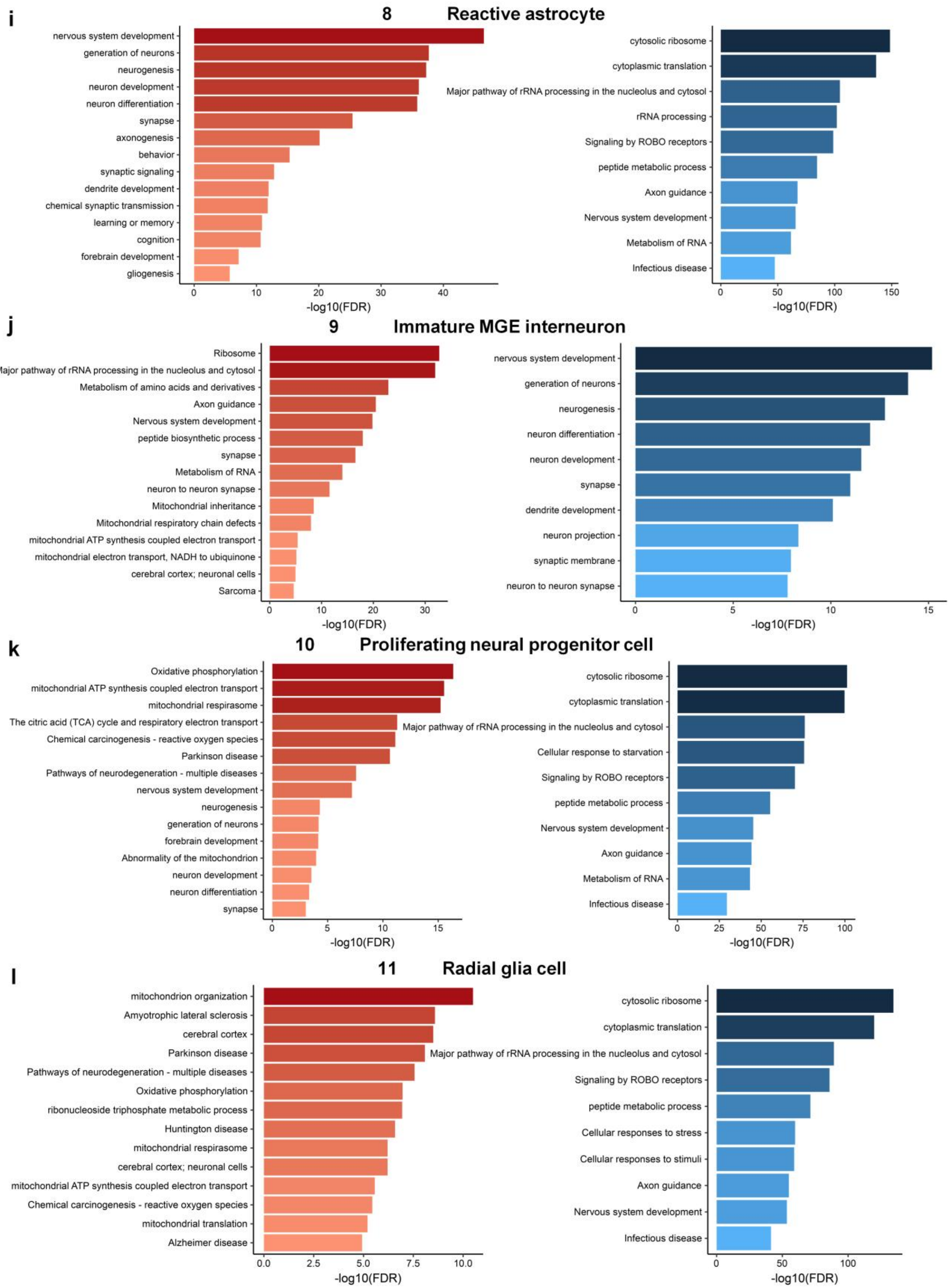

i
8 Reactive astrocyte
nervous system development
generation of neurons
neurogenesis
neuron development
neuron differentiation
synapse
axonogenesis
behavior
synaptic signaling
dendrite development
chemical synaptic transmission
learning or memory
cognition
forebrain development
gliogenesis
0 10 20 30 40
-log10(FDR)
cytosolic ribosome
cytoplasmic translation
Major pathway of rRNA processing in the nucleolus and cytosol
rRNA processing
Signaling by ROBO receptors
peptide metabolic process
Axon guidance
Nervous system development
Metabolism of RNA
Infectious disease
0 50 100 150
-log10(FDR)
j
9 Immature MGE interneuron
Ribosome
Major pathway of rRNA processing in the nucleolus and cytosol
Metabolism of amino acids and derivatives
Axon guidance
Nervous system development
peptide biosynthetic process
synapse
Metabolism of RNA
neuron to neuron synapse
Mitochondrial inheritance
Mitochondrial respiratory chain defects
mitochondrial ATP synthesis coupled electron transport
mitochondrial electron transport, NADH to ubiquinone
cerebral cortex; neuronal cells
Sarcoma
0 10 20 30
-log10(FDR)
nervous system development
generation of neurons
neurogenesis
neuron differentiation
neuron development
synapse
dendrite development
neuron projection
synaptic membrane
neuron to neuron synapse
0 5 10 15
-log10(FDR)
k
10 Proliferating neural progenitor cell
Oxidative phosphorylation
mitochondrial ATP synthesis coupled electron transport
mitochondrial respirasome
The citric acid (TCA) cycle and respiratory electron transport
Chemical carcinogenesis - reactive oxygen species
Parkinson disease
Pathways of neurodegeneration - multiple diseases
nervous system development
neurogenesis
generation of neurons
forebrain development
Abnormality of the mitochondrion
neuron development
neuron differentiation
synapse
0 5 10 15
-log10(FDR)
cytosolic ribosome
cytoplasmic translation
Major pathway of rRNA processing in the nucleolus and cytosol
Cellular response to starvation
Signaling by ROBO receptors
peptide metabolic process
Nervous system development
Axon guidance
Metabolism of RNA
Infectious disease
0 25 50 75 100
-log10(FDR)
l
11 Radial glia cell
mitochondrion organization
Amyotrophic lateral sclerosis
cerebral cortex
Parkinson disease
Pathways of neurodegeneration - multiple diseases
Oxidative phosphorylation
ribonucleoside triphosphate metabolic process
Huntington disease
mitochondrial respirasome
cerebral cortex; neuronal cells
mitochondrial ATP synthesis coupled electron transport
Chemical carcinogenesis - reactive oxygen species
mitochondrial translation
Alzheimer disease
0.0 2.5 5.0 7.5 10.0
-log10(FDR)
cytosolic ribosome
cytoplasmic translation
Major pathway of rRNA processing in the nucleolus and cytosol
Signaling by ROBO receptors
peptide metabolic process
Cellular responses to stress
Cellular responses to stimuli
Axon guidance
Nervous system development
Infectious disease
0 50 100
-log10(FDR)

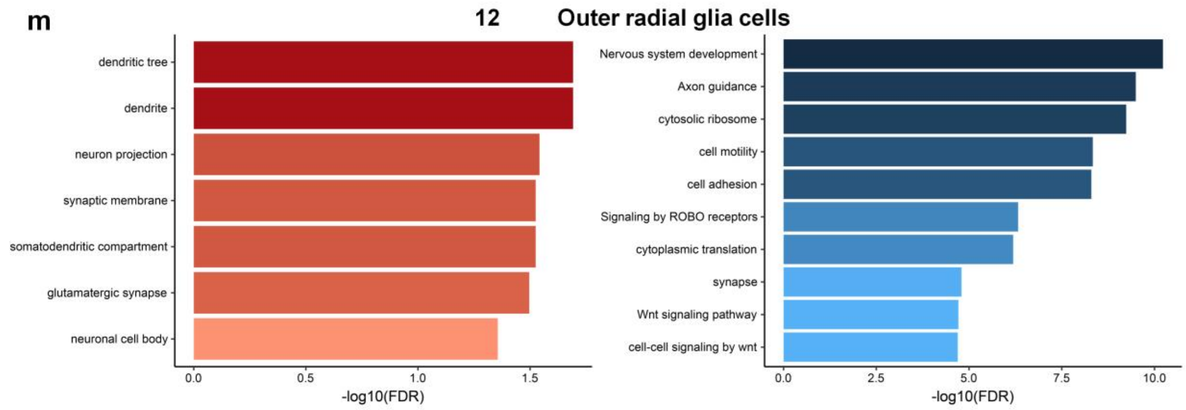


**Extended Data Figure 11. Gene ontology analyses highlight the main biological processes represented in each cell cluster.** For each cell cluster's DEGs from the single-cell transcriptomes, red bars represent GO terms are associated with upregulated genes (adjusted *P*-value < 0.05, log2 fold-change > 0.25), and blue bars represent those are associated with downregulated genes (adjusted *P*-value < 0.05, log2 fold-change < - 0.25).

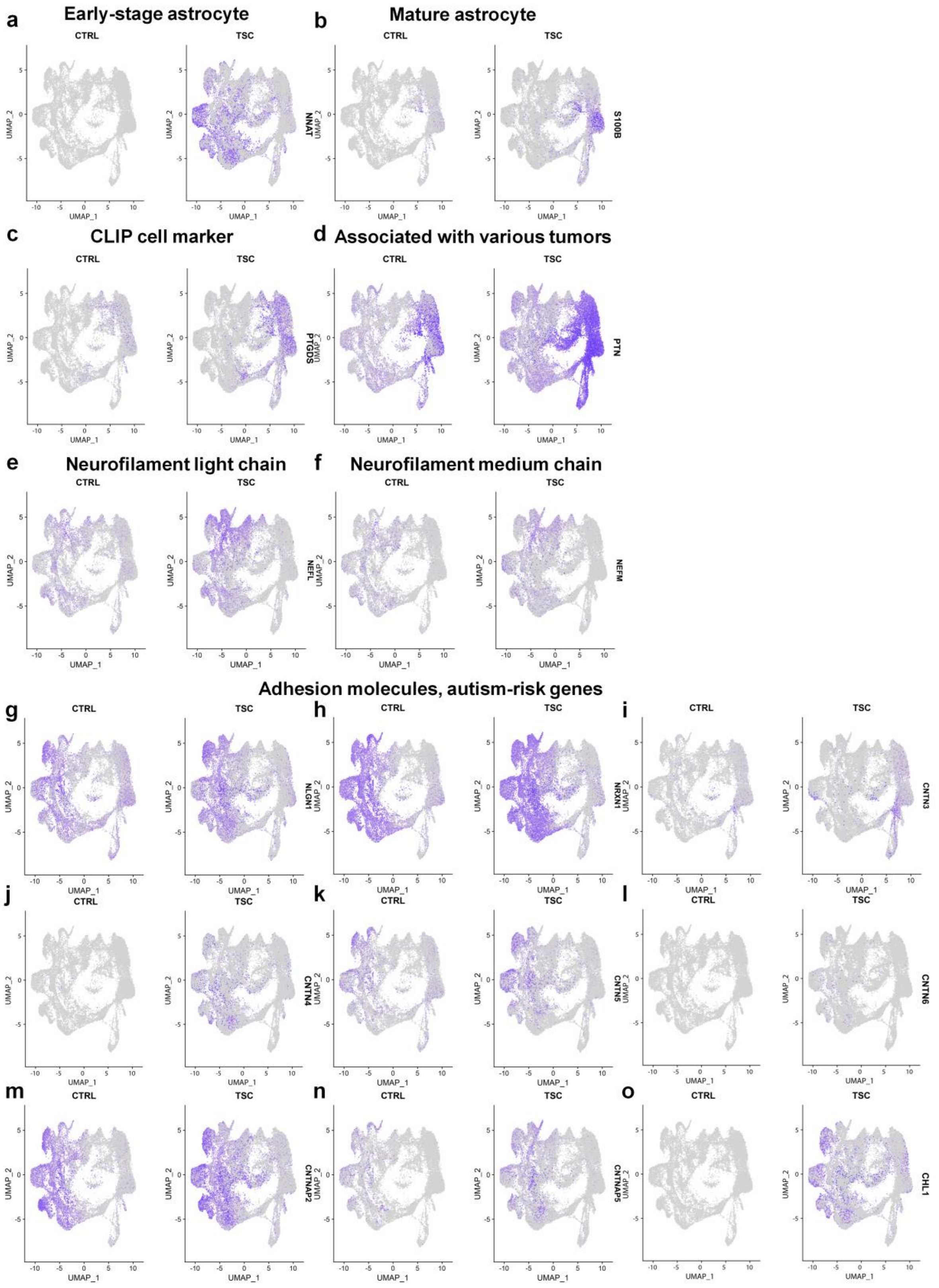
a
Early-stage astrocyte
b
Mature astrocyte
CTRL
TSC
NNAT
S100B
c
CLIP cell marker
d
Associated with various tumors
PTGDS
PTN
e
Neurofilament light chain
f
Neurofilament medium chain
NEFL
NEFM
Adhesion molecules, autism-risk genes
g
h
i
NLGN1
NRXN1
CNTN3
j
k
l
CNTN4
CNTN5
CNTN6
m
n
o
CNTNAP2
CNTNAP5
CHL1
UMAP_1
UMAP_2

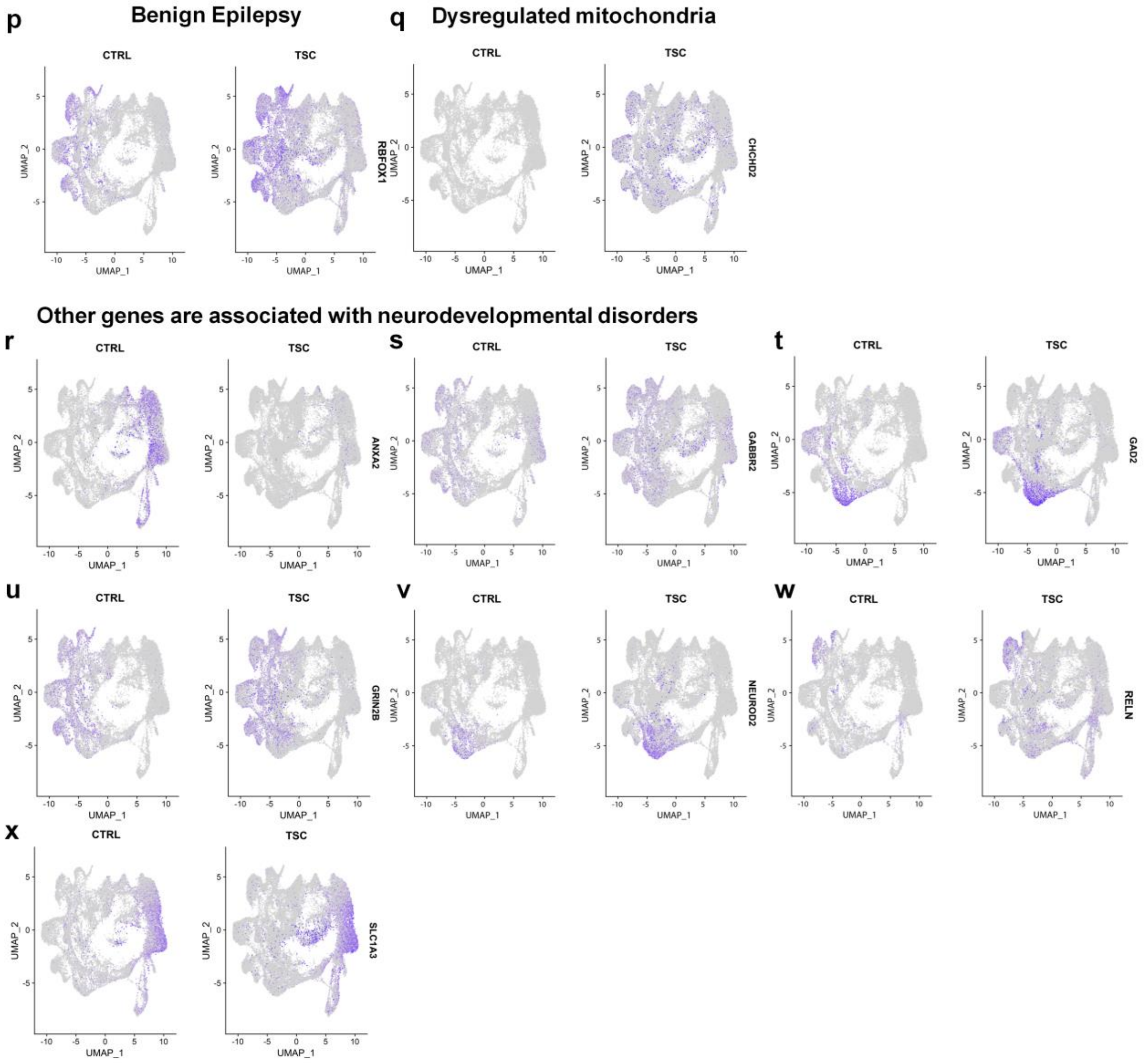


**Extended Data Figure 12. Genes associated with neurodevelopmental disorders are altered in TSC organoids.** UMAP plots show the expression of genes associated with neurodevelopmental disorders in both control and TSC organoids, including *NNAT*, *S100B*, *PTGDS*, *PTN*, *NEFL*, *NEFM*, *NLGN1*, *NRXN1*, *CNTN3*, *CNTN4*, *CNTN5*, *CNTN6*, *CNTNAP2*, *CNTNAP5*, *RBFOX1*, *CHCHD2*, *NEFL*, *ANXA2*, *GABBR2*, *GAD2*, *GRIN2B*, *NEUROD2*, *RELN*, and *SLC1A3*.

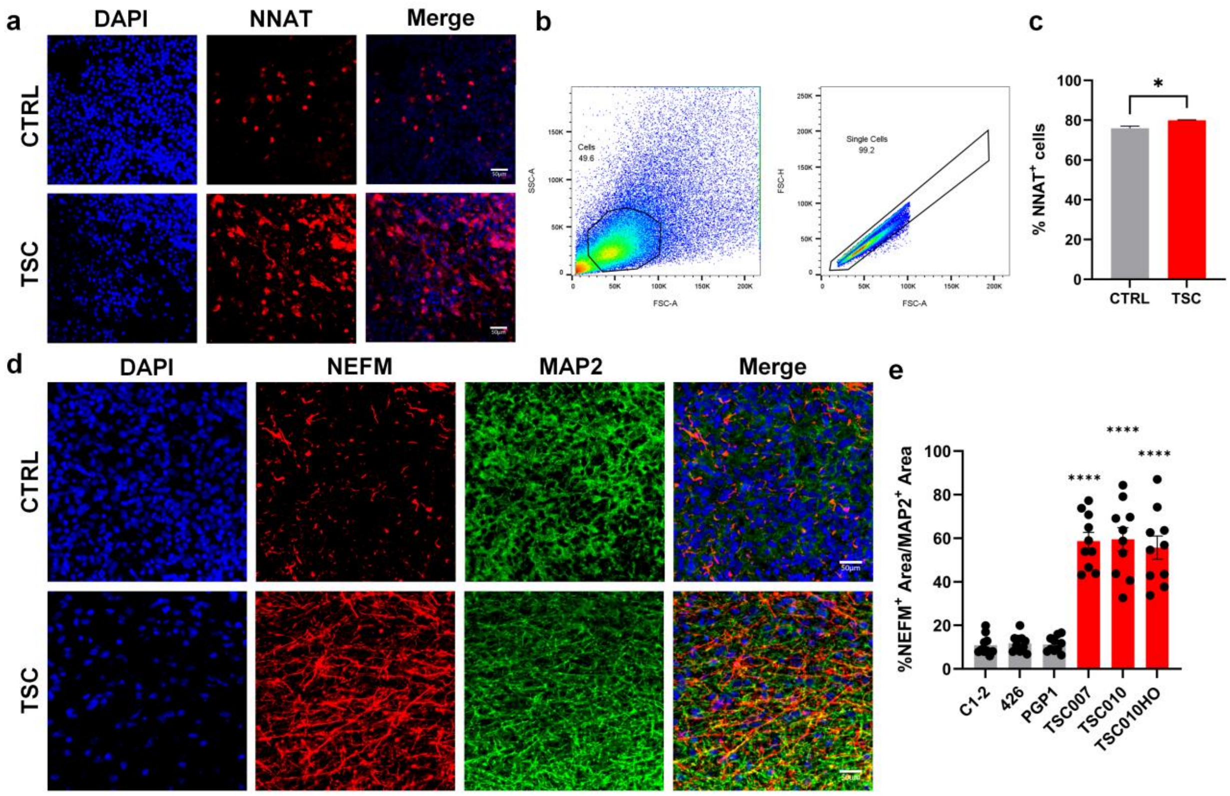


**Extended Data Figure 13. *TSC2* variants enhance early-stage astrocyte and NEFM formation.** **(a)** Shown are representative images of $NNAT^+$, an early-stage astrocytes marker, in both control and TSC organoids at Day 98, scale bars, 50 μm. **(b)** Gating strategy for quantification of the proportion of $NNAT^+$ cells by flow cytometry. Single cells, live cells were identified as depicted above. **(c)** Flow cytometry analysis of $NNAT^+$ staining indicates an increased proportion of $NNAT^+$ cells in TSC organoids compared to controls. Data are presented as mean ± s.e.m. (Unpaired t-test, *$P < 0.05$, n = 3 CTRL, 3 TSC). **(d-e)** Elevated NEFM level in TSC organoids. Shown are representative images (d) and quantification (e) of the percentage of $NEFM^+$ area relative to $MAP2^+$ area in control and TSC organoids. Data are presented as mean ± s.e.m. (n = 10 for each line from three independent experiments, ****$P < 0.0001$, one-way ANOVA), scale bars, 50 μm.

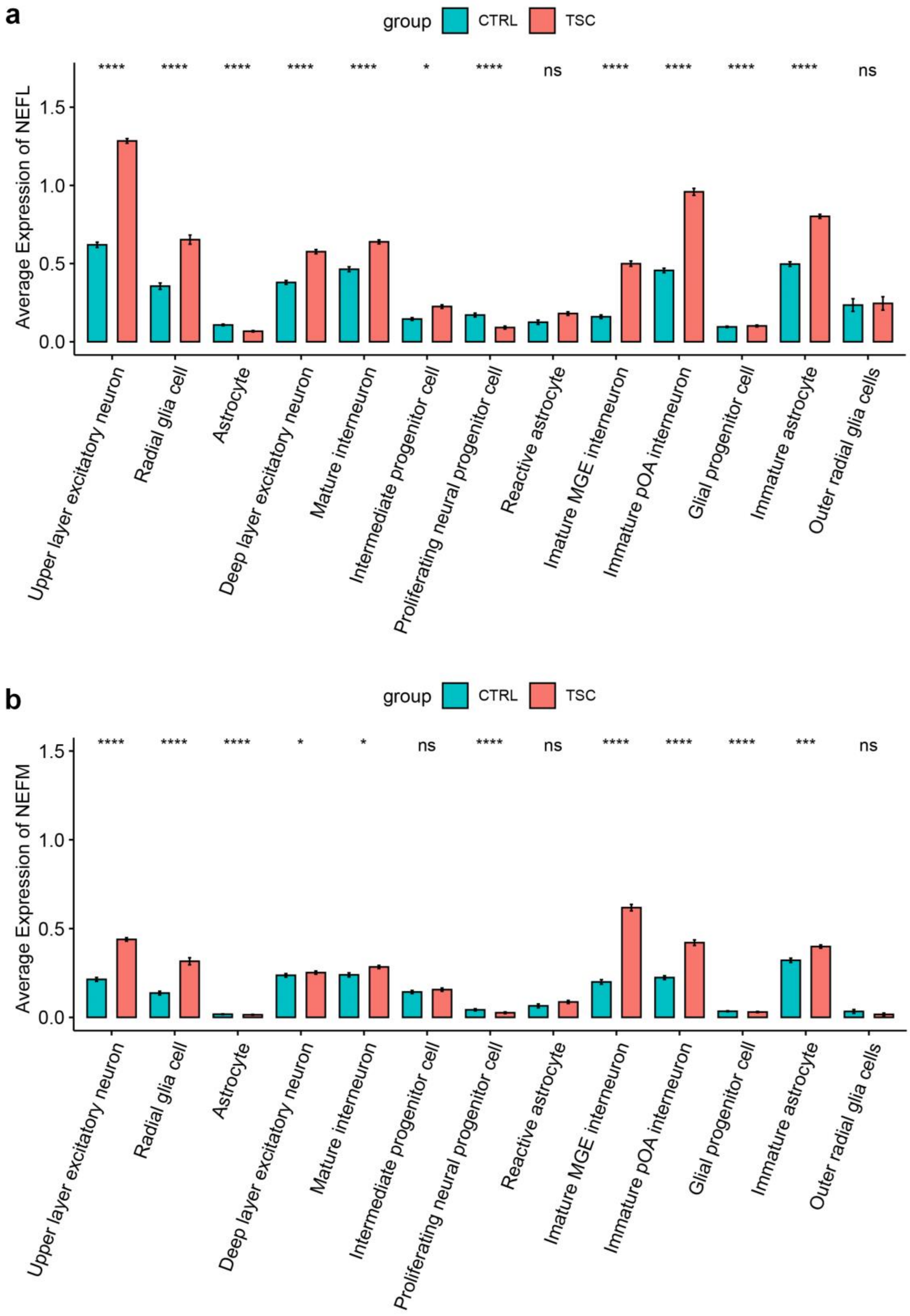


**Extended Data Figure 14. *TSC2* variants increase *NEFL* and *NEFM* in various types of cells.** **(a)** Average expression of NEFL. **(b)** Average expression of NEFM.

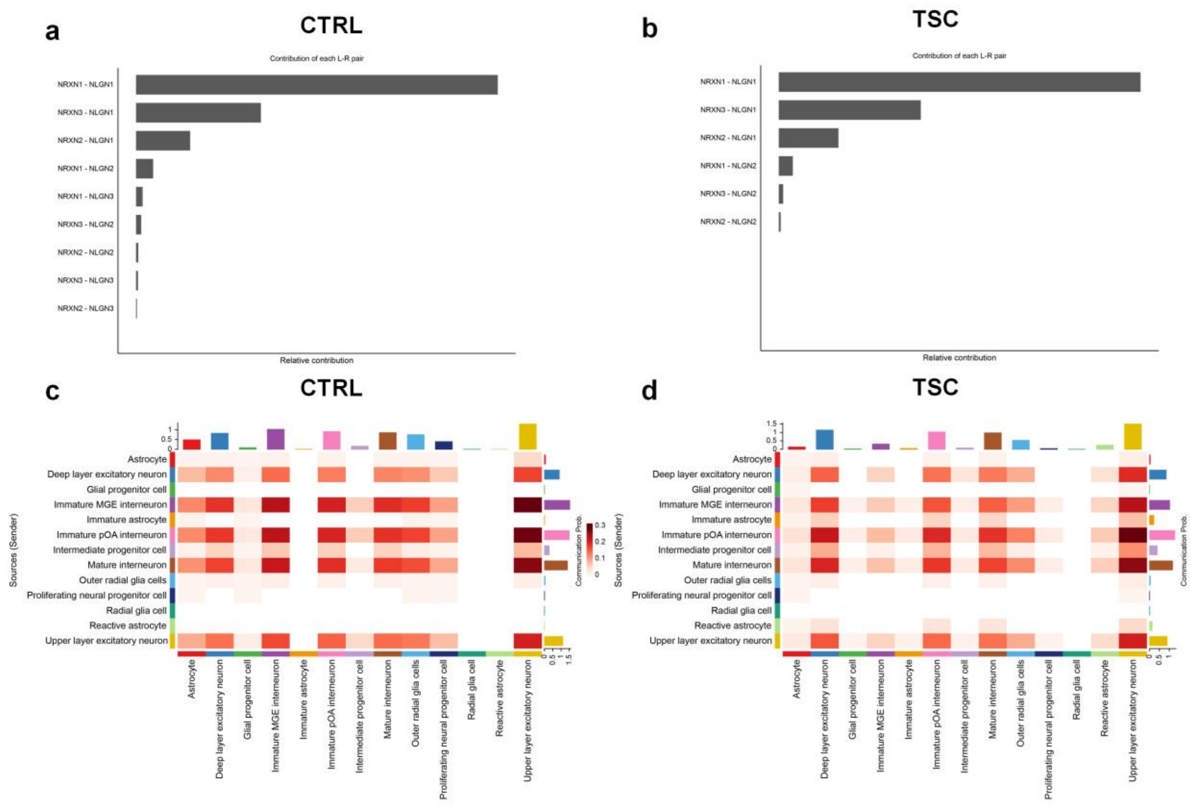


**Extended Data Figure 15. TSC organoids exhibit altered NLGN-NRXN signaling network for cell-cell communication. (a-b)** Calculate the contribution of each ligand-receptor pair within the overall NLGN-NRXN signaling network and visualize the cell-cell communication mediated by individual ligand-receptor pairs in control (a) and TSC organoids (b). **(c-d)** Heatmaps display the inferred NLGN-NRXN signaling network in control (c) and TSC (d) organoids, with color intensity between cell clusters indicating the strength of the interaction, as analyzed using CellChat.

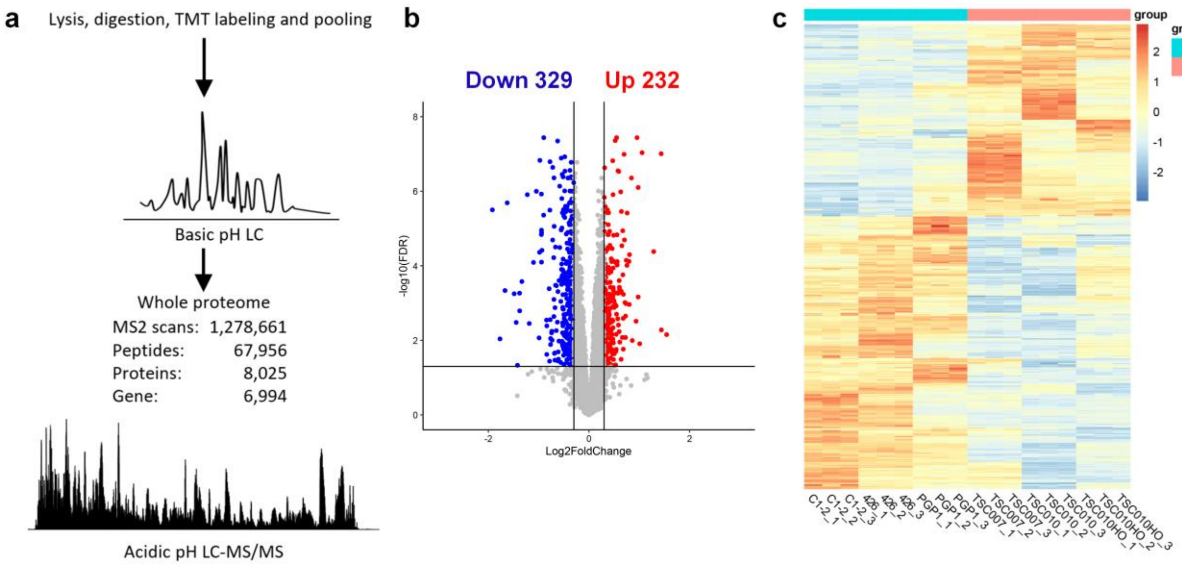


**Extended Data Figure 16. Proteomic analyses of the synaptosomes from control and TSC organoids. (a)** Proteomic workflow of 18-plex TMT-LC/LC-MS/MS. The study included three healthy controls and three TSC lines, each with three biological replicates. **(b)** Volcano plot of differentially expressed proteins (DEPs). Red and blue dots indicate 232 upregulated proteins and 329 downregulated proteins, respectively (*P*-value < 0.05, absolute log2 fold-change > 0.3). Grey dots represent proteins with no significant differential expression. **(c)** A heatmap is generated from the DEPs (*P*-value < 0.05, absolute log2 fold-change > 0.3) of TSC organoids compared to the controls. Colors indicate normalized protein expression levels.
